# Astrocytic Encoding of Threat in the Basolateral Amygdala

**DOI:** 10.64898/2026.01.07.697973

**Authors:** Ossama Ghenissa, Mathias Guayasamin, Sarah Peyrard, Ciaran Murphy-Royal

**Affiliations:** Département de Neurosciences, Université de Montréal, Montréal, Québec, Canada; Centre de Recherche du Centre Hospitalier de l’Université de Montréal (CRCHUM), Montréal, Québec, Canada

## Abstract

The basolateral amygdala (BLA) has long been implicated in threat detection and the generation of anxiety states. While previous experiments have demonstrated the important role of BLA principal neurons in driving anxiety-related behaviors, population-level recordings suggest that principal neurons encode broad exploratory states rather than anxiety *per se*. This discrepancy questions whether anxiety is indeed represented within the BLA, or if the BLA reflects a broader representation of behavioral states. Here, using simultaneous *in vivo* calcium recordings in both astrocytes and principal neurons in the BLA we find that in contrast to neurons, astrocytic activity provides a stable and scalable representation of threat-induced anxiety across an array of behavioral tasks. We find that the magnitude of astrocyte response to threatening stimuli is modulated by the individual anxiety levels of animals, and that exploration of anxiogenic environment can be decoded across multiple tasks using astrocyte activity alone. Our results shed light on a specialized encoding property of BLA astrocytes and establish these cells as key computational elements of threat processing in anxiety circuits.

## Introduction

Anxiety, defined as the anticipation of future threat^1^, is an evolutionarily conserved emotional state that promotes survival by enhancing vigilance and avoidance behaviors. While adaptive in nature, excess or misguided anxiety is a debilitating condition that can greatly impair one’s quality of life^1,2^. Anxiety disorders are among the most prevalent psychiatric disorders affecting around 300 million people worldwide^4^ and with a lifetime prevalence estimated ranging between 10 to 30%^5–7^. Among the many brain regions found to be implicated in the generation of anxiety states^8–12^, the basolateral amygdala (BLA) has emerged as a key integrator of emotional and behavioral responses associated with threat detection and anxiety-related behaviors^13–15^. The BLA has been shown to receive convergent anxiety-related inputs^16–20^ and has been proposed to serve as a critical node in fear circuits enabling rapid execution of behavioral change^21^. Beyond its strategic positioning within the anxiety network, direct manipulations of BLA activity or its projections to downstream targets have been shown to bidirectionally modulate anxiety-related behaviors^22–24^, making it a primary locus to understand anxiety processing.

Recent studies have highlighted a role for astrocytes in brain computation, regulating neural output from local synaptic tuning to modulation of global network dynamics^25–27^. Due to their diverse expression of receptors that are tuned to local neuronal circuits, their strategic positioning between synapses and blood vessels, and their ability to release neuromodulatory molecules (gliotransmitters), astrocytes have been demonstrated to integrate a multitude of signals in the extracellular milieu to fine-tune neuronal activity in a context-dependent manner^28^. At the network level, this astrocyte modulation of neuronal activity has been proposed to facilitate the selection of context-appropriate behavioral responses^29^, and is supported by an increasing body of literature demonstrating a role for astrocytes in behavioral tuning^30–41^. In the BLA, investigations of astrocyte function have mostly focused on fear conditioning, with astrocyte integrity important for long-term memory^42,43^. Despite this, it remains unknown whether astrocytes are actively implicated in distinct BLA-dependent processes including real-time encoding of emotional states and anxiety-like behavior.

In this study, we used *in vivo* calcium imaging in freely behaving mice to track the activity of both astrocytes and excitatory (principal) neurons in the BLA across an array of anxiogenic and threatening behavioral tasks. We report that astrocytes, but not principal neurons, provide a stable representation of anxiety-like states in spatial-as well as novelty-induced anxiety tasks. We find that the magnitude of astrocytic activity is linked to the anxiety levels of individual animals, and that exploration of anxiogenic environments can be readily decoded from one task to another using astrocyte calcium activity alone. Finally, we reveal that BLA astrocyte calcium activity is dynamically modulated by learned threat, showing rapid adaptation to auditory cued-fear conditioning. Together, these results reveal a specialized encoding property of astrocytes within the BLA and reveal that astrocyte calcium activity is a robust anxiety state indicator that is engaged in innately anxiogenic environments and by learned threat.

## Results

### Basolateral Amygdala Astrocytes Respond to Anxiogenic Environments

To investigate the activity of BLA astrocytes in anxiety-like states, we virally expressed GCaMP8f specifically in BLA astrocytes and recorded astrocyte calcium (Ca^2+^) activity in freely moving mice *via* fiber photometry across an array of anxiety-related tasks (**Fig. 1A** and **Figure S1A**). First, we verified that our approach using viral expression of GCaMP8f under the GfaABC1D promoter, was highly specific to BLA astrocytes (**Figure S1B-D**). We then evaluated astrocyte Ca^2+^ dynamics in BLA astrocytes during the exploration of prototypical anxiogenic environments using field-standard assays, starting with the elevated plus maze (EPM) a commonly used behavioral assay to assess anxiety-related behaviors (**Fig. 1B**). Behavioral analysis showed that mice spent significantly more time in the closed arms compared to the center, with least amount of time spent in the open arms (**Fig. 1C**). Analysis of astrocyte activity dynamics revealed robust increases in Ca^2+^ activity during exploration of the aversive zones of the maze (**Fig. 1B** and **Video S1**) with successively increasing calcium activity from closed arms to center zone and to open arms (**Fig. 1D**). Remarkably, these discrete increases in astrocyte Ca^2+^ activity were time-locked to behavioral transitions in the maze with increases in Ca^2+^ fluorescence during exits from closed arms into open arms and decreases upon re-entry into closed arms (**Fig. 1E**).

**Figure 1.**
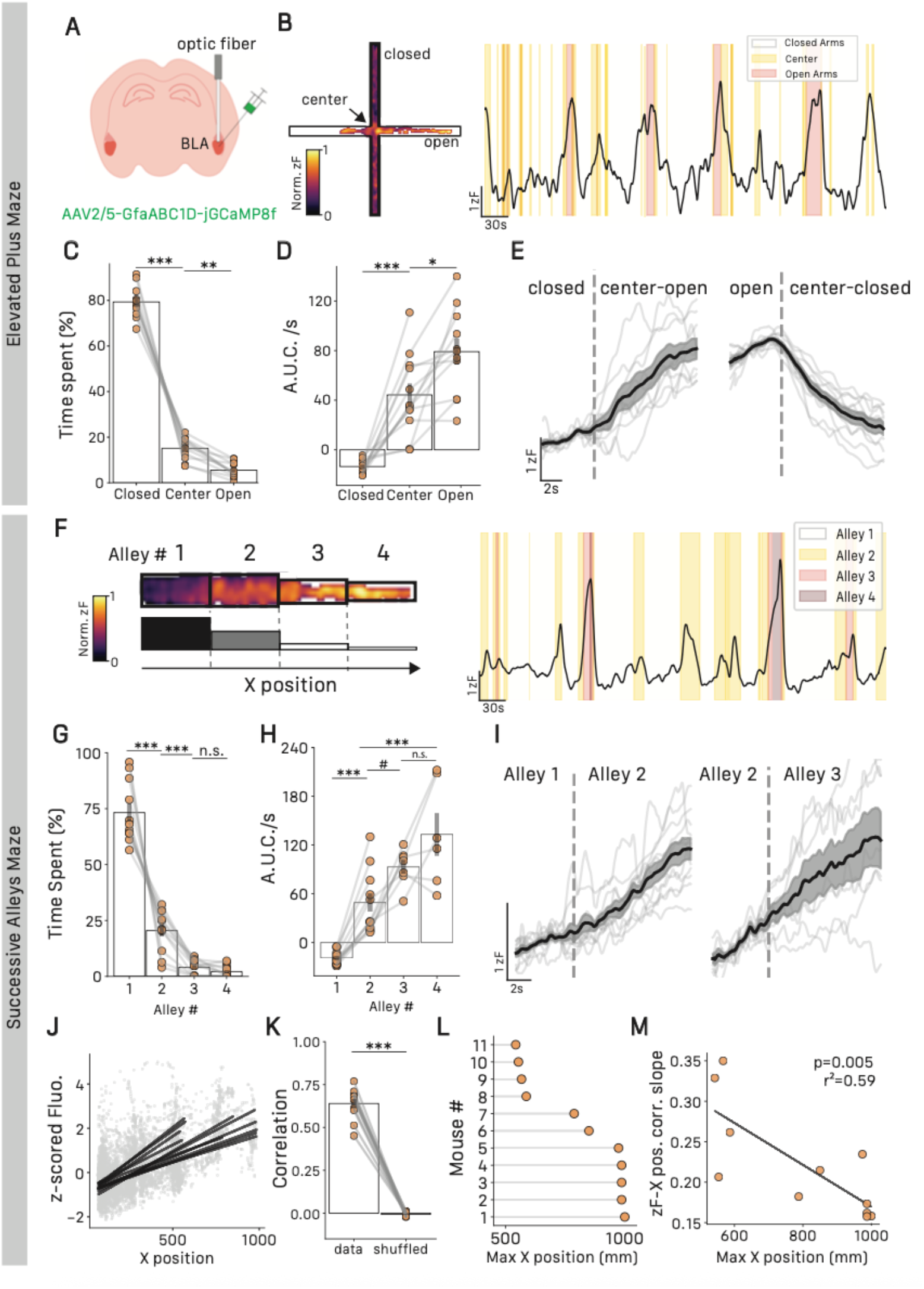
BLA Astrocytes respond to anxiogenic exploration. **(A)** Schematic of viral strategy and optic fiber implantation in the BLA. **(B)** Representative heat map (left) and trace (right) of astrocyte Ca2+ activity in the EPM. **(C)** Time spent in EPM compartments, closed = −79.35±2.13, center = 15.12±1.45, open = 5.51±0.98, repeated measure ANOVA F_2,20_ = 421.89, p<0.001, Tukey HSD closed vs. center p<0.001, center vs. open p=0.43, n=11 mice. **(D)** Area under the curve per second in EPM compartments, closed = −13.97±1.65, center = 48.23±9.15, open = 83±10.94, repeated measure ANOVA F_2,18_ = 42.9, p<0.001, Tukey HSD closed vs. center p<0.001, center vs. open p=0.014, n=11 mice. **(E)** Average z-scored fluorescence traces of astrocyte activity during closed arms exits (left) and entries (right), n=11, gray traces represent individual averages, n=11 mice. **(F)** Representative heat map (left) and trace (right) of astrocyte Ca2+ activity in the SAM. **(G)** Time spent in the SAM compartments, alley 1 = 72.97±4.12, alley 2 = 20.42±2.85, alley 3 = 4.04±1, alley 4 = 2.15±0.8, repeated measure ANOVA F_3,30_ = 122.4, p<0.001, Tukey HSD alley 1 vs. 2 p<0.001, alley 2 vs. 3 p<0.001, alley 3 vs. 4 p=0.95, n=11 mice. **(H)** Area under the curve per second in SAM compartments, alley 1 = −19.06±2.55, alley 2 = 49.15±11.73, alley 3 = 93.1±8.6, alley 4 = 133.11±26.56, repeated measure ANOVA F_3,15_ = 24.86, p<0.001, Tukey HSD alley 1 vs. 2 p<0.001, alley 2 vs. 3 p = 0.075, alley 2 vs. 3 p = 0.21, alley 2 vs. 4 p < 0.001, n=11 mice. **(I)** Average z-scored fluorescence traces of astrocyte activity during transitions from alley 1 to 2(left) and 2 to 3 (right), n=11, gray traces represent individual averages, n=11 mice. **(J)** Astrocyte z-scored fluorescence through mouse position in the SAM, black lines indicate individual regression lines, n=11 mice. **(K)** Coefficients from z-scored fluo. – mouse position against coefficients of a shuffled dataset, data = 0.63±0.02, shuffled = 0.027±0.003, paired T-test, T=21.99, p<0.001, n=11 mice. **(L)** Maximum X position reached by animals throughout testing, n=11 mice. **(M)** Correlation between slopes of z-scored fluorescence – mouse position correlation and maximum X position reached in the SAM, r^2^=0.59, p=0.005, n=11 mice. All results are represented as mean ± SEM. Graphical significance levels are *p<0.05; **p<0.01, and ***p<0.001.

Further behavioral analysis of exploratory behavior within the EPM revealed that all mice showed a preference for a specific closed arm (**Figure S2A,B**). We excluded that this preference was due to cues in the testing environment, as group-level preference between both arms remained equal (**Figure S2C**) however, we found that the preferred closed arm was often the first closed arm visited during testing (**Figure S2D**). We then set out to determine whether this subtle behavioral preference could be represented within BLA astrocyte activity. Remarkably, we found that BLA astrocyte Ca^2+^ activity was lowest in the preferred closed arm and increased during exploration of the non-preferred closed arm (**Figure S2E**), revealing a striking sensitivity of astrocyte Ca^2+^ signal with behavior.

While the EPM is a widely used measure of anxiety-like behavior, this behavioral apparatus is somewhat limited due to the binary nature of exploration of open versus closed arms. To understand the sensitivity of BLA astrocytes to more subtle changes in anxiogenic environments we tested the same cohort of mice in the successive alley maze (SAM). This linear maze consists of 4 successive compartments, ordered from the least aversive (Alley 1; black color floor and high black walls), to the most aversive compartment (Alley 4; narrow white color floor and no walls) (**Fig. 1F**). We hypothesized that astrocyte Ca^2+^ should scale with threat and exhibit highest activity in the most aversive compartment. The anxiogenic nature of this apparatus was quantified by measuring time spent in each compartment with mice spending most time in Alley 1 (similar to closed arm of EPM) and decreasing time spent in each subsequent compartment (**Fig. 1G**). Consistent with our hypothesis, astrocyte Ca^2+^ exhibited consistent increases in fluorescence scaled to the anxiogenic nature of the alleys, with a progressive increase from Alley 1 to 4 (**Fig. 1H**). Investigation of transitions between alleys revealed robust time-locked increases in activity when transitioning not only from the 1^st^ to the 2^nd^ but also from the 2^nd^ to the 3^rd^ alley (**Fig. 1I**).

To determine a relationship between astrocyte Ca^2+^ activity and anxiety states, we took advantage of the linearity of the SAM and considered the position of the mice along the main axis of the maze as a proxy of increased anxiogenic properties of the environment (referred to as X position, **see methods**). Using this metric, we found that position of the mice in the maze was highly correlated to fluorescence intensity of astrocyte Ca^2+^, with astrocyte activity linearly increasing with exploration of increasingly aversive alleys (**Fig. 1J,K**). Indeed, some mice explored the entire maze up to the last alley with peak activity in 4th alley, while other mice only ventured as far as the beginning of the 3rd alley, suggestive of individual variability in anxiety levels (**Fig. 1L**). Maximum exploration distance was used as an indicator of individual anxiety levels, as this measure strongly correlated with time spent exploring the anxiogenic alleys of the maze, a widely used metric of anxiety (**Figure S3A**). We leveraged this variability to assess whether individual anxiety levels (i.e. maximum distance explored in the maze) had an impact on astrocytic Ca^2+^ response. We found an inverse correlation between the slope of the signal-position relationship and the maximum exploration distance reached during the task, suggesting that mice with greater anxiety-like behavior exhibit stronger astrocytic Ca^2+^ activity at reduced distances along the alleys of the maze (**Fig. 1M**). Moreover, while the strength of the signal-position relationship evolves with anxiety levels, the magnitude of the maximal astrocyte Ca^2+^ activity remains equivalent between individuals (**Figure S3B**).

Consistent with the idea that astrocyte activity encodes anxiety across various behavioral tasks, we could apply the same linear analysis to the EPM task using the position of the mouse on the open arms axis (**Figure S3C,D**). Maximum exploration distance in the open arms of the EPM showed strong correlations between both time spent exploring in the EPM (**Figure S3E**) and maximum exploration distance in the SAM (**Figure S3F**), indicating that these measures of anxiety are stable within individuals across various tasks. Similar to what we found in the SAM, the position of the mice along the open arm in the EPM was strongly linked with astrocyte Ca^2+^ activity (**Figure S3G,H**). Consistently, more anxious mice exhibited lower maximum exploration distance and showed stronger increases in Ca^2+^ during open arm exploration (**Figure S3I**), while maximum astrocyte Ca^2+^ activity was equivalent between mice irrespective of individual anxiety levels (**Figure S3J**).

To eliminate confounding factors from our experiments we first investigated the possibility of habituation of the astrocyte Ca^2+^ activity during each behavioral task. Analysis epochs of the EPM and SAM trials were divided into two phases, 0-5 mins and 5-10 mins. Importantly, we found that astrocyte Ca^2+^ activity remained relatively stable through time, with no significant differences in z-scored fluorescence between these two epochs (**Figure S4A,B**). Additionally, we characterized BLA astrocyte Ca^2+^ dynamics in a novel but less anxiogenic environment, placing mice in an open field box (**Figure S5A**). We found that in this environment, mice spent more time in the periphery rather than the center zone of the maze (**Figure S5B**). When we quantified astrocyte Ca^2+^ activity by zone we observed a non-significant trend toward increased astrocyte activity during exploration of the center compared to the periphery (**Figure S5C,D**). Importantly, astrocyte Ca^2+^ dynamics showed no correlation with velocity, indicating that BLA astrocyte activity is not related to movement (**Figure S5E,F**).

Together, these results reveal a robust and stable relationship between BLA astrocyte Ca^2+^ activity and anxiety states, with increases in astrocyte Ca^2+^ activity proportional to the anxiogenic nature of the environment. Interestingly, the strength of this relationship was modulated by the anxiety levels of each individual, with higher astrocyte activity associated with reduced exploration of anxiogenic environments across EPM and SAM.

### BLA Astrocytes Respond to Novelty-Associated Threat, But Not Novelty in Familiar Environments

Despite our observations that BLA astrocyte Ca^2+^ dynamics remained stable throughout time, we wanted to explicitly test the role of novelty in gating astrocytic responses to potential threats as novel contexts have been documented to be anxiogenic in humans, rodents, and other organisms^44–47^. As such, rather than use EPM and SAM which rely on intrinsic anxiogenic properties of each maze, we adapted a behavioral task enabling recording of astrocyte Ca^2+^ activity during exploration of familiar and novel environments. To do this, mice were allowed 5 minutes to freely explore 2 arms of a Y-maze, commonly used to assess working memory^48^. Following this habituation phase, the 3^rd^ arm (identical but previously blocked) was made available for exploration during a testing phase (**Fig. 2A**). We first report that mice were able to discriminate between the novel and familiar arms, as they spent more time in the novel arm during testing (**Fig. 2B**). We then quantified astrocyte Ca^2+^ activity during the first 10 seconds of exploration of familiar versus novel arms and found significantly higher activity during exploration of the novel arm compared to the familiar arms (**Fig. 2C**). During the first exploration bout of the novel arm, BLA astrocyte Ca^2+^ activity showed a robust time-locked increase as the mice transitioned from the familiar to novel environment (**Fig. 2A,D**). Remarkably, this increase was only present for the first exploration of the novel arm, with subsequent exploration bouts exhibiting a phenotype similar to activity dynamics in familiar arms (**Fig. 2E**). These data suggest that astrocytic encoding of threat activity rapidly adapts once ambiguous environments have been assessed as non-threatening.

**Figure 2.**
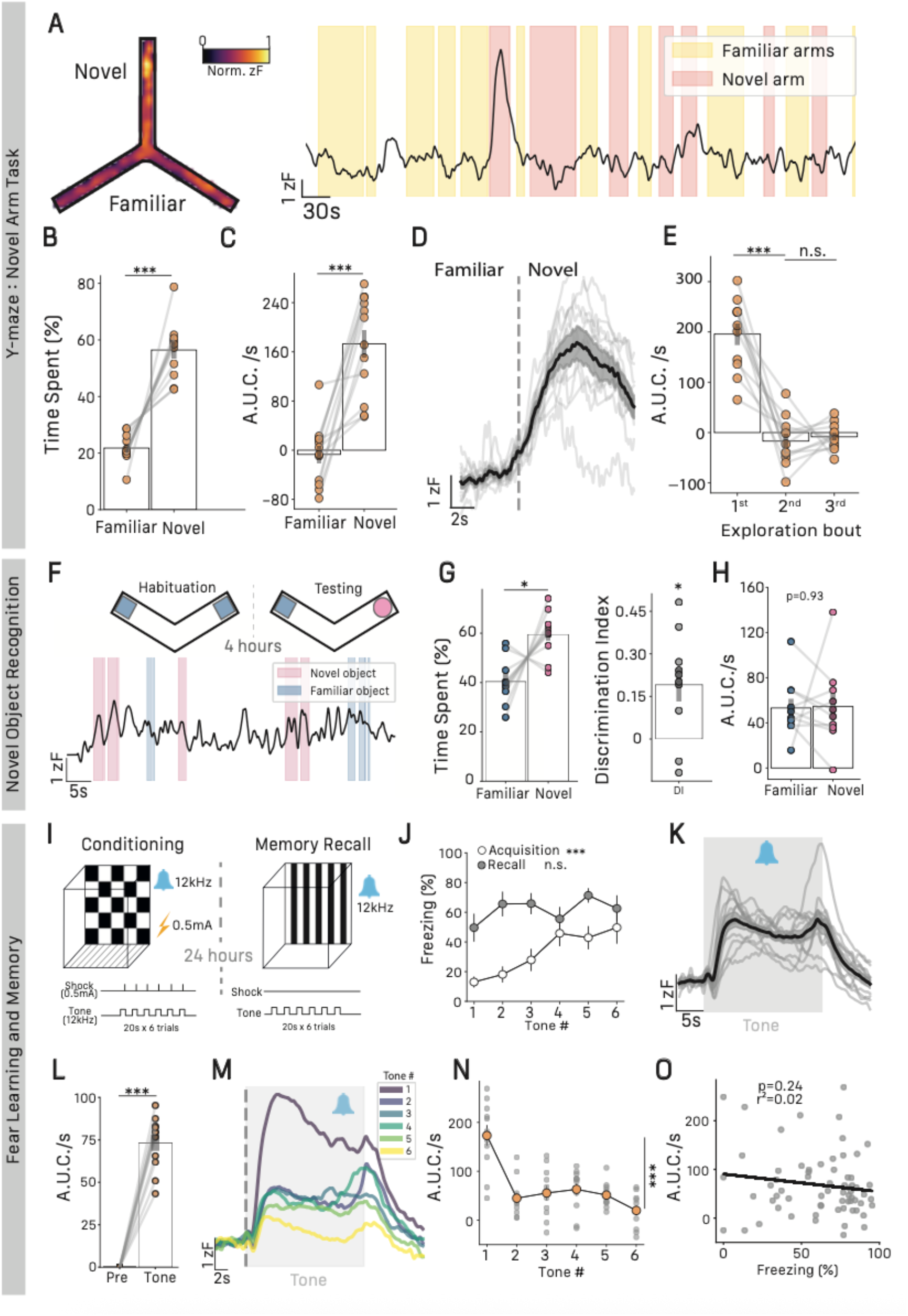
Rapid adaption of astrocyte activity to threat. **(A)** Representative heat map (left) and trace (right) of astrocyte-derived z-scored fluorescence in the Novel Arm Y-Maze task. **(B)** Percentage of time spent in familiar and novel arms of the Y-maze, familiar = 21.78±1.54, novel = 56.43±3.08, paired T-test, T=7.48, p<0.001, n=11 mice. **(C)** Area under the curve per second during 10 first seconds of exploration of familiar and novel arms of the Y-maze, familiar = −9.04±15.77, novel = 183.76±22.56, Wilcoxon Signed-Rank test, W≈0, p<0.001, n=11 mice. **(D)** Average z-scored fluorescence traces of astrocyte activity during first transition from familiar to novel arm, gray traces represent individual averages, n=11 mice. **(E)** Area under the curve per second for successive bouts of exploration of the novel arm. 1^st^ bout = 195.74±21.93, 2^nd^ bout = −16.92±14.64, 3^rd^ bout = −8.37±7.91, repeated measure ANOVA F_2,20_=49.89, p<0.001, n=11 mice. **(F)** Schematic of novel object recognition protocol (top) and representative z-scored fluorescence trace of 20 first seconds of object exploration (bottom). **(G)** Percentage of time spent exploring familiar and novel objects during 20 first seconds of exploration, familiar = 40.43±2.94, novel = 59.56±2.94, paired T-test, T=3.24, p=0.01, (left); Discrimination index of novel object exploration, DI=0.19±0.06, one-sided T-test, T=3.2, p=0.005 (right), n=9 mice. **(H)** Astrocyte area under the curve for the 10 first seconds of exploration of familiar and novel objects, familiar = 53.21±8.17, novel = 54.50±11.51, paired T-test T=0.08, p=0.93, n=9 mice. **(I)** Schematic of auditory fear conditioning paradigm. **(J)** Percentage of freezing through tone number for acquisition and recall phase, repeated measure ANOVA, effect of tone on freezing during acquisition, F_5,50_=6.64, p<0.001, effect of tone; Recall, repeated measure ANOVA, effect of tone on freezing during recall F_5,50_=1.41, p=0.26, n=11 mice. **(K)** Average z-scored fluorescence traces of astrocyte activity during tone presentation, gray traces represent individual averages, n=11 mice. **(L)** Area under the curve for pre-tone and tone presentation, pre-tone = 0±0, tone =73.2±4.81, paired T-test, T=15.21, p<0.0001, mice. **(M)** Average z-scored fluorescence traces of astrocytes activity through successive tone presentation during recall, gray traces represent individual averages, n=11 mice. **(N)** Area under the curve per sec of astrocyte activity through tone number during recall, repeated measure ANOVA, F_5,50_=17.02, p<0.001, n=11 mice. **(O)** Correlation between area under the curve and percentage time spent freezing during tone presentation, r^2^=0.02, p=0.24, n=11 mice. All results are represented as mean ± SEM. Graphical significance levels are *p<0.05; **p<0.01, and ***p<0.001.

To investigate potential differences in anxiety-related responses in this task, we quantified latency to enter the novel arm (**Figure S6A**). We found that latency to enter the novel arm was highly correlated to individual measures of maximum exploration distance in the EPM and SAM tests (**Figure S6B,C**), suggesting that latency to enter the novel arm is related to anxiety levels in individual mice. Interestingly, we found that latency to enter the novel arm was also correlated to the amplitude of the astrocytic Ca^2+^ response during the first exploration of the novel arm (**Figure S6D**), again suggestive of modulation of astrocyte response by individual anxiety levels.

To further elucidate the impact of novelty and determine whether astrocytes respond to any novel cue or only when there is a potential threat, we explored the response of BLA astrocyte Ca^2+^ dynamics during a novel object recognition test in a familiar environment (**Fig. 2F**). We show that mice were able to accurately discriminate between the novel and familiar object (**Fig. 2G**) however, in this task BLA astrocyte Ca^2+^ activity dynamics did not show important changes in activity during the initial exploration bouts of the novel object (**Fig. 2H**). Taken together, these data reveal that BLA astrocytes are highly tuned to respond to potentially threatening novel environments but not novelty in the absence of threat, i.e. in familiar environments.

### BLA Astrocytes Are Engaged During Amygdala-Dependent Fear Memory Recall

Having established a role for BLA astrocytes in encoding anxiety states, we next tested whether astrocytes also contribute to learned threat. To do this, we employed an amygdala-dependent auditory-cued fear conditioning paradigm. In brief, on the conditioning day mice were exposed to conditioned stimulus (auditory cue, 12kHz pure tone, 20s duration) that co-terminated with a foot shock (0.5mA, 2s duration). Twenty-four hours later, amygdala-dependent fear memory was tested by exposing mice to the conditioned stimulus in a novel environment (**Fig. 2I**). During fear acquisition, mice showed increased freezing behavior throughout tone-shock pairing that persisted twenty-four hours later during memory recall, indicative of fear learning and memory (**Fig. 2J**). Astrocyte Ca^2+^ activity exhibited dynamic responses to successive tone and footshock presentations during the acquisition phase, indicative of encoding of learned threat associated with this conditioning paradigm (**Figure S7A-C**). During fear memory recall, BLA astrocyte Ca^2+^ activity showed a persistent increase upon conditioned stimulus presentation (**Fig 2K,L**).

Interestingly, we found adaptation in the astrocytic Ca^2+^ signal which rapidly decreased upon successive tone presentations (**Fig 2M,N**). This decrease, however, was not related to changes in freezing behavior as we found no correlation between astrocyte Ca^2+^ activity and freezing behavior during tone presentation (**Fig 2O**). Nevertheless, this rapid adaptation of the astrocyte response in this learned behavioral task is consistent with our findings of innate behavior in the Y-maze, in that astrocyte Ca^2+^ response decreases when threat is removed from the cue or environment.

### Astrocytic Activity is Uncoupled from Local BLA Pyramidal Neuron Activity

As BLA principal neurons have been implicated in many aspects of anxiety-related behavior, we set out to understand the potential relationship between astrocyte activity dynamics and local excitatory neuron activity. We used dual-color photometry to simultaneously record astrocyte Ca^2+^ activity using GCaMP8f and neuronal Ca^2+^ activity by expressing jRGECO1a in CaMKII-expressing cells (**Fig. 3A**). Using this approach, we were first able to reproduce our initial findings in the EPM (**Fig. 3B,C**) showing again that astrocyte Ca^2+^ activity increases with exploration of center and open arms of the maze (**Fig. 3D**). In contrast, investigation of neuronal Ca^2+^ dynamics failed to reveal a robust distinction between EPM compartments (**Fig. 3B,E**). Additionally, we did not observe changes in neuronal Ca^2+^ activity time-locked to transitions between those compartments (**Figure S8A,B**). Second, using the SAM we found opposite modulation of astrocytic and neuronal Ca^2+^ activity between maze compartments (**Fig. 3F**). Time spent in each compartment was consistent with the previous cohort (**Fig. 3G**) however, while astrocytic Ca^2+^ activity increased with exploration of anxiogenic alleys (i.e. 2^nd^, 3^rd^, 4^th^) neuronal Ca^2+^ activity decreased (**Fig. 3H,I; Figure S8C,D**). Third, in the Y-maze novel arm task, mice once again were able to discriminate between novel and familiar arms (**Fig. 3J**) and in this task both astrocytic (**Fig. 3K**) and neuronal (**Fig. 3L**) Ca^2+^ activity responded with robust, transient increases during the first exploration of the novel arm (**Figure S8E,F**). Consistently, neuronal Ca^2+^ activity showed the same adaptive properties as for astrocytes in this task, with the Ca^2+^ activity dynamics returning to that of familiar arm exploration at 2^nd^ and subsequent exploration bouts (**Figure S8G**). In the novel object recognition task, in which mice were readily able to discriminate novel objects from familiar (**Fig. 3M**), astrocyte Ca^2+^ activity did not reliably relate to discrimination between familiar and novel objects (**Fig. 3N**) whereas neurons actively responded to novelty (**Fig. 3O**) revealing divergence in neuronal and glial responses to this neutral valence learning and memory task.

**Figure 3.**
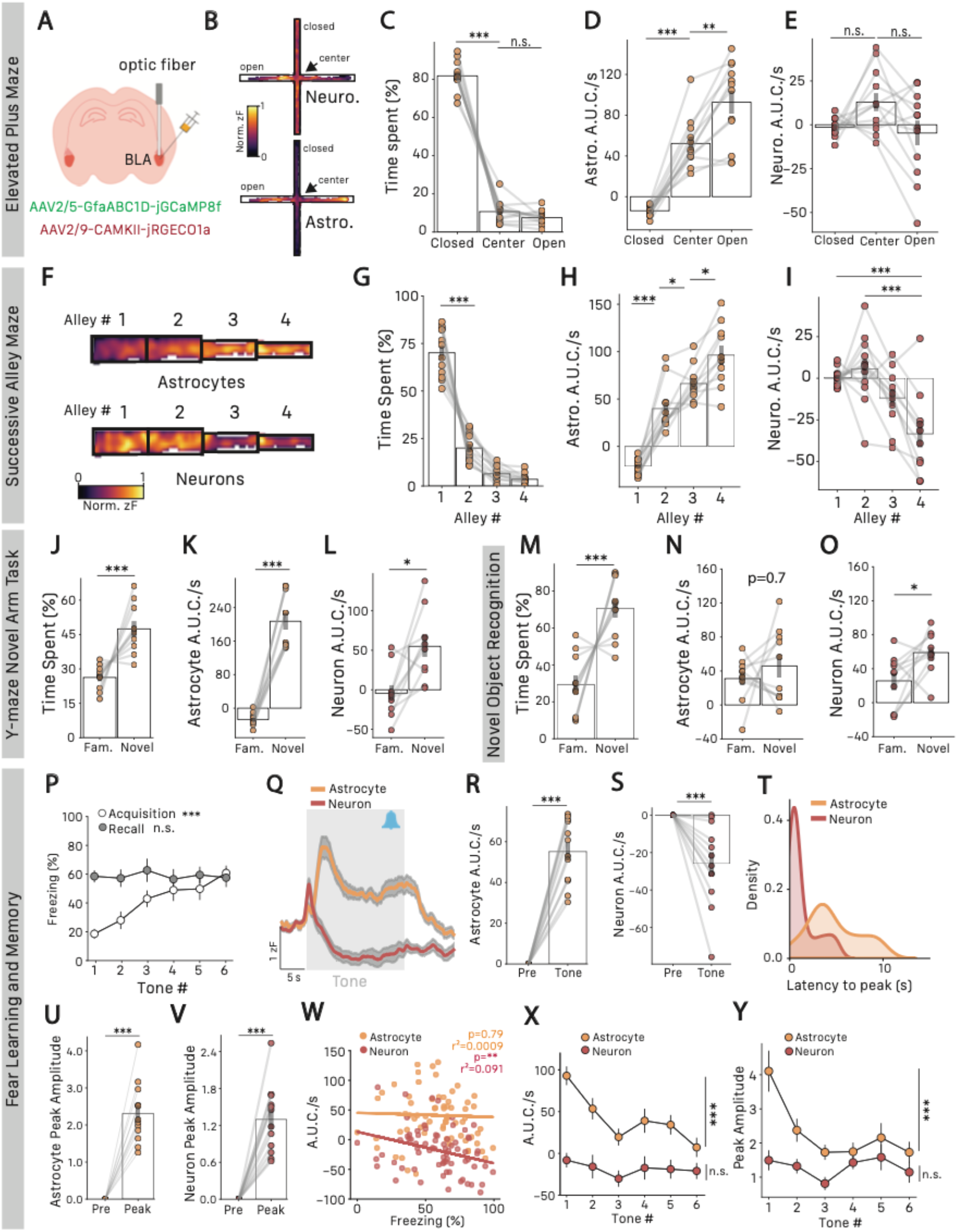
Astrocyte activity is uncoupled to local neuron activity. **(A)** Schematic of dual color photometry viral strategy and optic fiber implantation in the BLA. **(B)** Representative heat map of neuronal (top) and astrocytic (bottom) Ca2+ activity in the EPM. **(C)** Percentage of time spent in EPM compartments, closed = −81.75±2.25, center = 10±1.69, open = 7.56±1.06, repeated measure ANOVA F_2,22_ = 387.94, p<0.001, Tukey HSD closed vs. center p<0.001, center vs. open p=0.42, n=12 mice. **(D)** Astrocytes z-scored area under the curve per second in EPM compartments, closed = −13.84±1.51, center = 52.25±7.1, open = 92.76±11.61, repeated measure ANOVA F_2,22_ = 68.79, p<0.001, Tukey HSD closed vs. center p<0.001, center vs. open p=0.002, n=12 mice. **(E)** Neurons z-scored area under the curve per second in EPM compartments, closed = −1.43±1.5, center = 12.92±5.15, open = −4.69±7.18, repeated measure ANOVA F_2,22_ =3.26, p=0.057, Tukey HSD closed vs. center p=0.13, center vs. open p=0.055, n=12 mice. **(F)** Representative heat map of neuronal (top) and astrocytic (bottom) Ca2+ activity in the SAM. **(G)** Time spent in the SAM compartments, alley 1 = 70.1±3.6, alley 2 = 19.98±2.12, alley 3 = 6.42±1.15, alley 4 = 3.54±0.82, repeated measure ANOVA F_3,32_ = 147.06, p<0.001, Tukey HSD alley 1 vs. 2 p<0.001, alley 2 vs. 3 p<0.001, alley 3 vs. 4 p=0.79, n=12 mice. **(H)** Astrocytes z-scored area under the curve per second in SAM compartments, alley 1 = −21.05±2.39, alley 2 = 39.89±66.99, alley 3 = 66.3±5.51, alley 4 = 96.36±10.63, repeated measure ANOVA F_3,27_ = 88.06, p<0.001, Tukey HSD alley 1 vs. 2 p<0.001, alley 2 vs. 3 p = 0.036, alley 3 vs. 4 p = 0.019, n=12 mice. **(I)** Neurons z-scored area under the curve per second in SAM compartments, alley 1 = 0.71±1.5, alley 2 = 5.73±5.89, alley 3 = −11.84±5.2, alley 4 = −33.4±8.23, repeated measure ANOVA F_3,27_ = 12.03, p<0.001, Tukey HSD alley 1 vs. 2 p=0.9, alley 2 vs. 3 p =0.11, alley 3 vs. 4 p = 0.0504, alley 1vs.4 p<0.001, alley 2 vs. 4 p<0.001, n=12 mice. **(J)** Percentage of time spent in familiar and novel arms of the Y-maze, familiar = 26.27±3.43, novel = 47.44±1.71, paired T-test, T=4.11, p=0.002, n=11 mice. **(K)** Astrocytes area under the curve per second during 10 first seconds of exploration of familiar and novel arms of the Y-maze, familiar = −27.27±4.96, novel = 207.24±20.44, paired T-test, T=10.82, p<0.001, n=11 mice. **(L)** Neuron area under the curve per second during 10 first seconds of exploration of familiar and novel arms of the Y-maze, familiar = −4.33±10.06, novel = 59.94±13.85, paired T-test, T=3.62, p=0.005, n=11 mice. **(M)** Percentage of time spent exploring familiar and novel object, familiar = 29.35±5.15, novel = 70.65±5.15, paired T-test, T=5.66, p<0.001, n=10 mice. **(N)** Astrocytes area under the curve for the 10 first seconds of exploration of familiar and novel objects, familiar = 31.01±8.33, novel = 45.9±13.71, paired T-test T=23, p=0.7, n=10 mice. **(O)** Neurons area under the curve for the 10 first seconds of exploration of familiar and novel objects, familiar = 25.741±9.89, novel = 58.99±7.72, paired T-test T=2.64, p=0.016, n=10 mice. **(P)** Freezing behavior through tone presentation during habituation and recall. Acquisition, repeated measure ANOVA, effect of tone F_5,55_=8.13, p<0.001; Recall, repeated measure ANOVA, effect of tone F_5,55_=0.2, p=0.96, n=11 mice. **(Q)** Average z-scored fluorescence traces of astrocytes and neuronal activity during recall phase tone presentation, n=11 mice. **(R)** Astrocytes area under the curve per second for pre-tone vs. tone presentation, pre-tone=0±0, tone=55.15±4.53, paired T-test, T=12.17, p<0.001, n=11 mice. **(S)** Neurons area under the curve per second for pre-tone vs. tone presentation, pre-tone=0±0, tone=-25.96±6.39, paired T-test, T=4,06, p<0.001, n=11 mice. **(T)** Kernel density estimation of latency to peak distributions for astrocyte and neuronal activity, Kolmogorov Smirnov test, p<0.001. **(U)** Astrocyte peak fluorescence during tone presentation, pre = 0±0, peak = 2.3±0.23, paired T-test, T=9.67, p<0.001, n=11 mice. **(V)** Neuronal peak fluorescence during tone presentation, pre = 0±0, peak =1.29±0.16, paired T-test, T=8.09, p<0.001, n=11 mice. **(W)** Astrocytes and neuronal area under the curve per second correlation to time spent freezing during tone presentation, linear regression, neurons r^2^=0.091, p<0.009, astrocytes r^2^=0.0009, p=0.79, n=11 mice. **(X)** Astrocytes and neuronal area under the curve per second through tone presentation during recall, repeated measure ANOVA, effect of tone on astrocytes F_5,55_=2.73, p=0.02, effects of tone on neurons F_5,55_=1.21, p=0.31, n=11 mice. **(Y)** Astrocytes and neuronal peak amplitude through tone presentation during recall, repeated measure ANOVA, effect of tone on astrocytes F_5,55_=7.88, p<0.001, effects of tone on neurons F_5,55_=1.33, p=0.28, n=11 mice. All results are represented as mean ± SEM. Graphical significance levels were *p<0.05; **p<0.01, and ***p<0.001.

Finally, to compare astrocyte and neuronal Ca^2+^ dynamics during amygdala-dependent learning and memory we employed auditory-cued fear conditioning. During fear acquisition, mice exhibited increased freezing with each tone-shock pairing on acquisition day as well as robust freezing to tone alone on recall day (**Fig. 3P**). Indeed, both astrocytes and neurons showed robust calcium responses to tone and foot-shock during conditioning (**Figure S8H-M**). During memory recall, we report dynamic cue-associated changes in both astrocyte and neuronal Ca^2+^ activity (**Fig. 3Q**). As anticipated, astrocyte Ca^2+^ activity displayed a persistent increase during tone exposure (**Fig. 3R**). Quantification of neuronal activity during the entire tone presentation, revealed a persistent decrease below baseline (**Fig. 3S**) however, we observed a peak response in both astrocytic and neuronal activity following tone onset, with neuronal activity more tightly tuned to tone onset (**Fig. 3T**). Analyses of peak responses to tone onset revealed a strong increase in both astrocytic (**Fig. 3U**) and neuronal calcium activity (**Fig. 3V**). We correlated these dynamic changes in astrocytic and neuronal signals with behavior to reveal that the decrease in the neuronal activity observed with A.U.C. analysis correlated with percentage time spent freezing to the conditioned auditory cue, this relationship was absent for astrocytic activity (**Fig. 3W**). This relationship between behavior and activity was absent for both astrocytes and neurons using the peak amplitude analysis (**Figure S8N,O**). Finally, to determine within trial adaptation of threat encoding we compared astrocytic and neuronal activity across the six exposures to the conditioned stimulus. We again observed rapid adaptation of the astrocyte signal to the auditory cue using both A.U.C. (**Fig. 3X**) or peak amplitude (**Fig. 3Y**) measures. This adaptation did not occur in the neuronal response (**Fig. 3X,Y**), suggestive of distinct encoding of information between astrocytes and neurons. These data suggest that astrocytic encoding of threat is highly dynamic showing rapid within-trial adaptation whereas adaptation of neuronal activity, which directly modulates behavior, occurs on a much slower timescale and aligns with active learning and memory processes associated with fear conditioning paradigms^49^

Overall, we demonstrated that during exploration of anxiogenic environments, astrocyte Ca^2+^ activity is independent of local excitatory neurons, with astrocytes exhibiting coherent and stable anxiety-related activity throughout tasks while neurons exhibit noisier, less stable changes in Ca^2+^ activity during anxiogenic exploration. Nevertheless, with learned threat neuronal activity was a more reliable readout of behavior with neuronal Ca^2+^ activity encoding memory in both novel object recognition and cued fear memory recall. These data highlight diverging roles of glia and neurons in brain computation and support the idea that astrocytes reliably encode anxiety and learned threat.

### Decoding Anxiogenic Context from Astrocyte Activity

If indeed astrocytes encode anxiety states, astrocyte Ca^2+^ activity should exhibit a robust signature across a wide range of anxiogenic contexts. To test this in an unbiased manner, we trained a logistic regression decoder for each mouse to classify exploration of the anxiogenic compartments in the SAM, using only z-scored astrocyte Ca^2+^ activity (**see Methods, Fig. 4A**). We then applied this SAM-trained decoder to predict exploration of anxiogenic compartments in the EPM task from astrocyte Ca^2+^ data alone (**Fig. 4B**). The decoder, trained on an entirely different task, significantly outperformed a control decoder trained on shuffled-label datasets, achieving 82 ±2.5% accuracy in prediction of whether the mouse was in open arms or center/closed arms of the maze based on astrocyte calcium dynamics (**Fig. 4C,D**). We further confirmed the robustness of this approach by training a new decoder on EPM data and testing its performance on SAM data (**Fig. 4E,F**). We found similar results, with on average 79 ±3.1% of accuracy for the decoder trained on true data compared to 50 ±0.4% for the shuffled control decoder (**Fig. 4G,H**). Both decoders consistently outperformed their respective controls on a variety of other performance metrics such as receiver operating characteristic AUC, f1-score, recall, and precision (**Figure S9A-H**). These results demonstrate that BLA astrocytes display a reliable Ca^2+^ activity signature during the exploration of anxiogenic versus anxiolytic contexts, regardless of the task.

**Figure 4.**
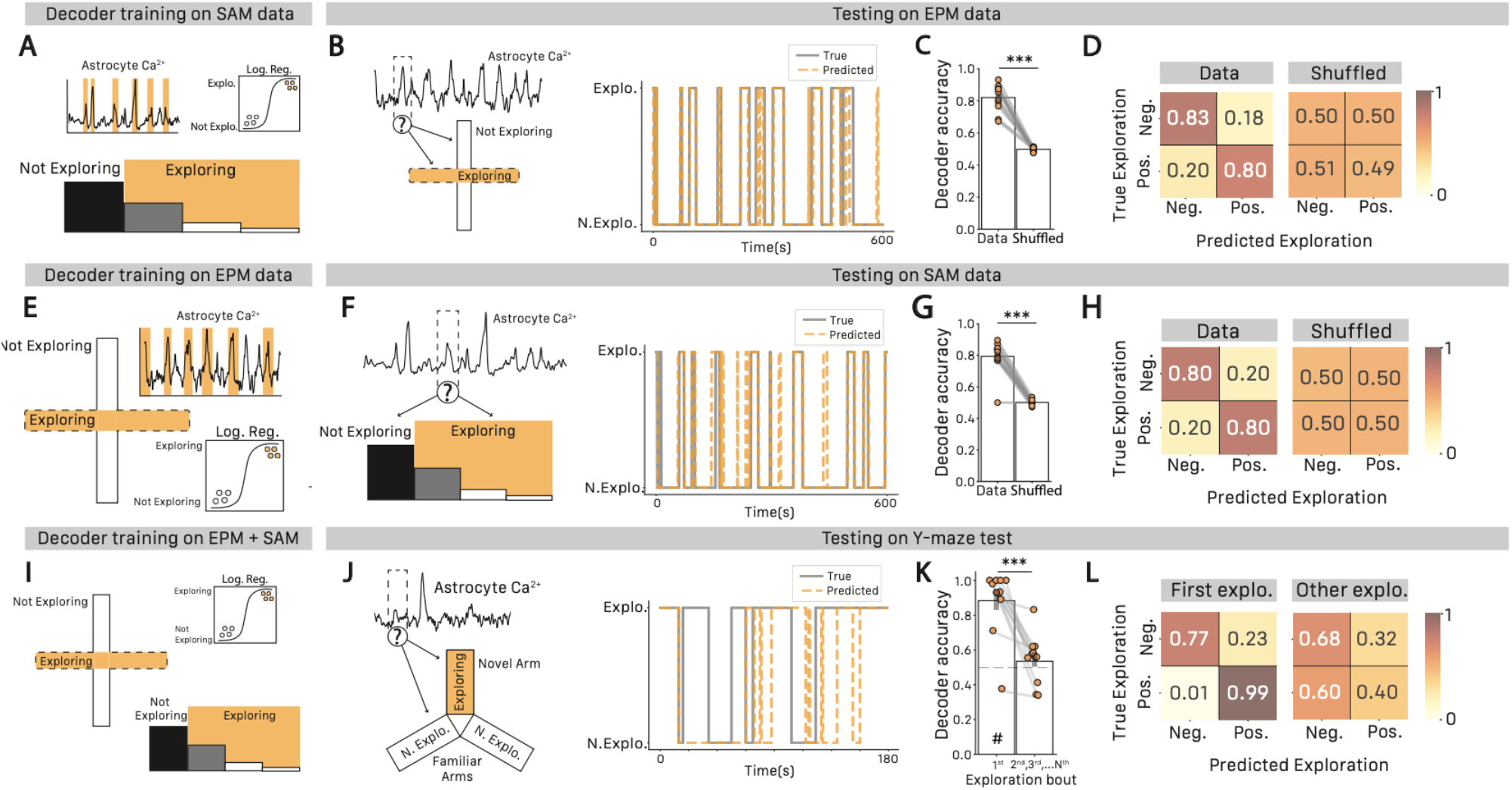
Decoding anxiogenic context from astrocyte calcium activity. **(A)** Schematic of logistic regression decoder training on SAM behavioral and astrocytic Ca^2+^ data. **(B)** Schematic of logistic regression decoder testing on EPM data (left), Example decoding of exploration of anxiogenic compartment in EPM from SAM-trained decoder (right). **(C)** Decoding accuracy from true data and shuffled labels trained decoders. True data trained decoder = 0.82±0.025, shuffled data trained decoder = 0.497±0.004, paired T-test, T=12.63, p<0.001, n=11 mice. **(D)** Average confusion matrix from decoders trained on true data (left) and shuffled data (right), n=11 mice. **(E)** Schematic of logistic regression decoder training on EPM behavioral and astrocytic Ca^2+^ data. **(F)** Schematic of logistic regression decoder testing on SAM data (left), Example decoding of exploration of anxiogenic compartment in SAM from EPM-trained decoder (right). **(G)** Decoding accuracy from true data and shuffled labels trained decoders. True data trained decoder = 0.793±0.031, shuffled data trained decoder = 0.501±0.004, paired T-test, T=9.016, p<0.001, n=11 mice. **(H)** Average confusion matrix from decoders trained on true data (left) and shuffled data (right), n=11 mice. **(I)** Schematic of logistic regression decoder training on EPM and SAM behavioral and astrocytic Ca^2+^ data. **(J)** Schematic of logistic regression decoder testing on Y-Maze Novel arm task data (left), Example decoding of exploration of novel arm in Y-maze novel task from EPM+SAM-trained decoder (right). **(K)** Decoding accuracy from Y-maze decoding of first vs. subsequent exploration of novel and familiar arms. First exploration = 0.883±0.005, subsequent exploration = 0.535±0.043, paired T-test, T=6.64, p<0.001 – dashed lines indicate performance of shuffled label trained decoder. # indicates statistical significance against shuffled-labels trained decoder, n=11 mice. **(L)** Average confusion matrix from Y-maze decoding trained for first exploration (left) and subsequent exploration (right), n=11 mice. All results are represented as mean ± SEM. Graphical significance levels were *p<0.05; **p<0.01, and ***p<0.001. # indicates significance against shuffled labels decoder equivalent.

We next tested whether we could decode anxiety states from 2 distinct types of anxiety-related tasks. We trained a decoder on both EPM and SAM data and tested it on our Y-maze novel arm dataset, considering exploration of the novel arm as anxiogenic exploration (**Fig. 4I,J**). Decoding the first exploration of each arm achieved high performance, with 88 ±1% accuracy when decoding exploration of novel versus familiar arms (**Fig. 4K** and **Figure S9I-M**) and a strong decoding bias towards the positive class (99% of correctly decoded epochs in novel arm for 77% of correctly decoded epoch in the familiar arm; **Fig. 4I**). This decoder, however, was unable outperform a control decoder on any scoring metrics with subsequent exploration bouts following the first bout (**Fig. 4K,L** and **Figure S9N-R**). This difference in decoding performance between first and subsequent explorations aligns with our hypothesis that astrocyte Ca^2+^ rapidly adapts when threat is no longer present, and the decoder lacked a reliable Ca^2+^ signature.

Finally, we investigated whether a decoder trained on neuronal Ca^2+^ data would also achieve robust inter-task decoding properties. Using both neuronal Ca^2+^ and astrocyte Ca^2+^ activity data from our dual-color photometry experiments, we trained 2 separate decoders on either astrocytic or neuronal activity for each mouse. For the EPM-to-SAM and SAM-to-EPM configurations, the neuronal decoder performed slightly above chance levels for most of our scoring metrics (all but f1-score and precision for EPM-to-SAM, and precision for SAM-to-EPM; **Figure S10A-J**). The astrocytic decoder trained on the same mice consistently outperformed the neuronal decoder on all performance metrics, performing well above chance and significantly better than the decoder trained on the neuronal dataset (**Figure S10A-J**). In the EPM+SAM-to-Y-maze configuration, we found similar results regarding decoding of the first exploration bout with neuronal decoder performing above chance and astrocyte decoder performing above neurons for all performance metrics (**Figure S10K-O**), while performance remained at chance level for both neuronal and astrocytic decoder when looking at subsequent exploration bouts, as expected (**Figure S10P-T**).

Globally, these results indicate that BLA astrocytes provide a stable and robust representation of threat that can be reliably used to predict anxiety-related behaviors across multiple tasks and contexts.

## Discussion

Our study reveals a novel role for BLA astrocytes in encoding innate anxiety and learned threat. Using a longitudinal approach to assess BLA astrocyte and principal neuron activity in the same mice across an array of anxiety-inducing tasks, we show that astrocytes robustly respond to anxiogenic environments, whereas neurons show less stable representations of anxiogenic contexts. BLA astrocytes consistently exhibited increases in Ca^2+^ activity in the EPM and SAM (two spatial anxiety tasks) and in the Y-maze novel arm task (a novelty-related anxiety task) but did not respond in the novel-object recognition test, a salience-based task. Indeed, astrocyte activity was strongly associated with individual anxiety states with peak activity associated with maximum exploration of anxiogenic environments. Moreover, we find that astrocyte activity is highly dynamic with rapid adaptation following removal of threat from the environment or from a learned cue. Finally, we find that astrocytes provide a stable representation of anxiety states that enabled us to decode anxiogenic exploration across distinct tasks. Overall, these data suggest that BLA astrocytes encode threat-associated anxiety states and do so more reliably than local neuronal counterparts.

In the elevated plus maze, successive alley, and Y-maze novel arm tasks, behavior was consistent across tests with individual mice exhibiting higher anxiety levels in one task also showing heightened anxiety in the others. This behavioral consistency suggests the presence of stable, innate predispositions to anxiogenic stimuli, indicative of trait anxiety^50^. A key finding from our data is that the magnitude of astrocytic responses in anxiogenic contexts was positively associated with higher anxiety-like phenotypes, i.e. reduced exploration within each maze. This suggests that heightened astrocyte activity influences behavioral responses to potential threat and that differing astrocyte calcium dynamics between individuals might explain variability in trait anxiety. Supporting this idea, we observed a linear increase in astrocyte calcium activity along the track of the successive alley maze, consistent with the progressive increase in anxiogenic nature of this task. This relationship highlights the scalability of astrocytic responses to environmental threat levels. Indeed, we observed a similar linear increase in activity along the open arms of the EPM which was somewhat surprising considering the uniformity of the environment across the arms. These independent confirmations suggest that BLA astrocyte calcium activity scales with threat and as such we observed lowest astrocyte Ca^2+^ activity occurring in the subjective safest zones i.e. preferred arm of EPM. We propose that this effect may reflect environmental cues with increasing distance from the enclosed center potentially amplifying perceived threat.

While previous experiments have firmly demonstrated the functional importance of BLA principal neurons in anxiety-related behaviors^17,23^, single-cell resolution *in vivo* imaging studies describe BLA principal neuron population activity as ensembles that represent global exploratory states rather than anxiety states *per se*^51,52^. This divergence raised the question of whether the BLA encodes anxiety directly, as observed in the ventral hippocampus^8,53^, or instead represents more global behavioral states. Recent evidence, however, demonstrated that projections to the BLA effectively relay anxiety-related information^16,54,55^, contrasting with the idea that anxiety is not represented within the BLA. Our results indicate that anxiety representation in principal neurons is highly variable and inconsistent across tasks, with astrocyte activity providing a reliable representation of anxiety. We did not observe any significant differences in neuronal activity between compartments in the EPM, found a decrease in activity throughout the SAM, and an increase in activity in the Y-maze novel arm task. Astrocytic activity, however, remained stable throughout all tasks, showing robust and scalable increases in activity when mice were confronted with anxiogenic stimuli. As such, our results may reconcile this discrepancy in the literature by suggesting a model in which astrocytes integrate anxiety-related information in the BLA.

Neuronal activity increased upon novel object exploration, consistent with previous findings suggesting that BLA neurons respond to saliency^56–58^, and showed an inverse relationship between activity during tone presentation and freezing, consistent with previous research on threat-related behaviors^59^. These learning and memory tasks are known to be encoded by discrete ensembles of neurons, forming what is known as an engram^60,61^. Astrocytes, however, did not respond to either exploration of the novel object or behavioral expression of conditioned fear but did respond to potential and learned threat. This suggests that astrocytes in the BLA are embedded within a distinctly specialized functional network outside of salience coding and fear memory expression. Our data align with other studies demonstrating a compartmentalized function of astrocytes including hippocampal astrocytes encoding reward location rather than acting as place cells^62^, astrocytes accumulating evidence of futility^63,64^, astrocytic gating of the effects of norepinephrine on synaptic efficacy^65^, and astrocytes closing critical periods of motor^66^ as well as visual^67^ plasticity. In this present work, we add to this functional specialisation of astrocytes revealing an enduring representation of anxiety states at the population level within the BLA.

In sum, our results reveal a unique astrocytic sensitivity to anxiety states that is uncoupled to the response of nearby neurons. This novel role for BLA astrocytes as specialized threat detectors in innate emotional processing and learned fear, positions BLA astrocytes as integral components of anxiety circuits. More generally, our work brings a new perspective to our understanding of the neural representation of anxiety states. As these data reveal a previously undescribed role of astrocytes in threat detection, our work could have broad implications for pathological anxiety disorders.

## Acknowledgements

We would like to thank all lab members for their input at all stages of this project. Ossama would specifically like to thank Meriam Zid for her precious input throughout the design and writing process. We thank Aurélie Cleret-Buhot (CRCHUM cellular imaging core) for microscopy training, the staff of the animal facility at the CRCHUM, Tamás Füzesi, Alexander Lohman and the CSM optogenetics core facility at the University of Calgary for training on fiber photometry approaches, and Neurophotonics summer school at Université Laval for imaging training.

## Author contributions

O.G. and C.M-R. designed the study. O.G. and M.G. carried out stereotaxic surgeries for viral injections and fiber implantations. O.G. carried out all experiments and analysis with help from M.G. for learning and memory experiments. S.P. maintained mouse colonies and provided expertise along those lines. O.G. and C.M-R. wrote the manuscript. All authors approved of the final version of the manuscript.

## Methods

### Animals

Both male and female C57BL/6J mice (Jax #000664) were used in this study. Mice were co-housed in groups of 3 to 5 with ad libitum food and water access. Housing rooms were kept on a 12:12 light-dark cycle and animals were always tested during light-cycle. All experiments were conducted in accordance with the guidelines for maintenance and care of animals of the Canadian Council on Animal Care and approved by the Institutional Committee for the Protection of Animals at the Center Hospitalier de l’Université de Montréal (protocol number CM20031CMRs).

### Stereotaxic surgeries and optic fiber implantation

5-6 weeks-old mice were subcutaneously injected carprofen (20mg/kg) before surgery. Mice were anesthetized with isofluorane (2.5% induction, 1.5% maintenance) and placed on a stereotaxic frame (Kopf Instruments) using non-rupture ear bars (Kopf instruments). Animal heads were shaved and eye-ointment (Systane) was applied. Bupivacaine/lidocaine (2 mg/kg) was locally applied before antero-posterior incision of the scalp at the midline. A small craniotomy was performed using a hand-held electric drill (Foredom). Viral vectors were unilaterally injected in BLA using a 10μL Hamilton neuro-syringe (Harvard Apparatus) at coordinates −1.4mm AP, ±3.2mm ML from bregma, and −4mm DV from surface of the brain at AP-ML coordinates. For all viral deliveries, a volume of 500nL was injected at a flow rate of 200nL/min with a microinjection syringe pump (UMP3T-2 World Precision Instruments). After injection was completed, the syringe was left in place for 7 minutes to allow for the fluids to settle, before slowly removing the needle.

Optic fibers were implanted during the same procedure. First, the skull was carved around the craniotomy for better adherence of the dental cement. The optic fibers (400 μm core, 0.66 NA, Doric Lenses) were mounted on a cannula holder (Doric Lenses) and slowly lowered at the injection site to −3.9 DV from brain surface. A first layer of C&B Metabond dental cement (Parkell) was applied around the cannula and left to dry for at least 5 mins. A second layer of dental cement (Teets ‘Cold Cure’, Diamond Springs) was finally applied to secure the implant. Mice were finally subcutaneously injected with 500μL of sterile saline and moved to a recovery room for at least 1h in a heated cabinet.

### Viral constructs

All viral vectors were packaged by the COVF viral vector core (RRID: SCR_016477). We used: AAV2/5-gfaABC1D-jGCaMP8f (lot 5347) at 5.1 x 10^12^ gc/mL, and 1:1 mix of AAV2/5-gfaABC1D-jGCaMP8f (lot 5347) 5.1 x 10^12^ gc/mL and AAV2/9-CAMKII-jRGECO1a (lot 4695) 5.1 x 10^12^ gc/mL.

### Behavioral assays

Before the start of experiments, mice were habituated to the testing room for 1 hour prior to handling by the experimenter for at least 3 days.

Mice were then habituated to the patch cord of the fiber photometry set up in their home cage for at least 2 sessions of 10 minutes. Experiments were completed in the following order: open field, elevated plus maze, successive alley maze, Y-maze test, Novel Object Recognition test. For the Astro-GCaMP8f group, auditory fear conditioning experiments were held on a separate cohort of mice. All behavioral apparatus were cleaned with 70% ethanol between each mouse unless stated otherwise.

#### Open field

The apparatus consisted of a 40×40×30 cm (length x width x height) white plexiglass box. Mice were placed at the center of the maze at the start of the experiment and left free to explore for 10 minutes.

#### Elevated Plus Maze

The maze consisted of 2 closed and 2 open arms of length 35cm and width 5.5cm, with a square center part of 5cm^2^, set up 75cm above the ground. At the start of the experiment, mice were placed at the center part of the maze and left free to explore for 10 minutes.

X position was computed as position of the mice along the open arm axis. The center of the maze was considered to be 0mm, and increased symmetrically throughout both open arms, with the extremity of both arms being considered as 375mm. Max. X position was considered as the maximum position reached by the mouse during the entire recording.

If mice fell from the EPM, overall testing data were kept but all related behavioral and photometry recordings were excluded from analysis (n=3 mice for Astro-gcamp8f group and n=2 mice for Astro-GCaMP8f+Neuro-jRGECO1a group).

#### Successive Alleys Maze

The maze consisted of 4 successive linear alleys of 25cm lengths. First alley was black, of width 8.5cm and wall height 25cm; second alley was grey of width 8.5cm and wall height 5cm; third alley was white, of width 3.5cm and wall height 0.8cm; fourth alley was white, of width 1.2cm and wall height 0.2cm. At the start of experiment, mice were placed at the beginning of the first alley, facing the wall, and were left free to explore for 10 mins.

X position was considered as the position of the mice along the main axis of the maze. The back wall of the first alley was considered to be 0mm while the extremity of the fourth alley was considered 1000mm. Max. X position was considered as the maximum position reached by the mouse during the entire recording.

#### Y-maze test

Apparatus consisted of a Y-maze with 3 arms of length 30cm and width 5.5cm. At the start of the experiment, only 2 arms of the maze were made available for 5 minutes of habituation. After 5 minutes, the experimenter waited for mice to get near the edge wall of one of the 2 available arms to begin the test phase. At the start of the test phase, the guillotine door preventing the access to the 3^rd^ arm was slowly removed and mice were then left free to explore until a total test time of 10 mins. Proportion of time spent in familiar vs. novel arm was computed on the first 3 minutes of exploration of the test phase.

#### Novel Object Recognition

The novel object recognition test protocol used here is a slightly modified version of *Da Cruz et Al*., *2020*. The tests were performed using our previously described Y-maze with 2 arms available. As the mice were already familiarized with the maze, the test did not include a habituation phase and consisted only of an acquisition and a test phase.

During the acquisition phase, 2 identical objects were placed at the extremities of the 2 arms of the maze, and mice were left free to familiarize with the objects for 9 mins. The test phase was held 4 hours later. During the test phase, a familiar object and a novel object (of around the same size but different in color and shape) were placed at the extremities of each arm. Mice were then left free to explore any object for 9 mins. The location of novel object on right or left arm was counterbalanced by random assignment of object location.

Exploration of an object was defined as behavioral bouts where mice were facing the object at a distance of maximum 0.5cm. Time spent exploring an object was defined as the proportion of time spent exploring the object during the first 20s of cumulative exploration of both objects. Discrimination index was computed as follows, with positive indices corresponding to more exploration of the novel object and negative indices indicating more exploration of the familiar object:

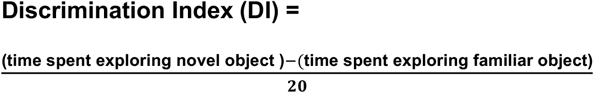

#### Auditory Cued Fear Conditioning

Auditory-cued fear conditioning tests were conducted in a Ugo Basile conditioning chamber with an electrified grid floor (Stoelting) of width 17×17×25cm (length x width x height). Testing consisted of a conditioning and a recall phase.

During conditioning (day 1), mice were put in the conditioning chamber in a first context (checkered walls cleaned with 70% ethanol). Mice were exposed to a tone (CS+; 12 kHz tone, 20 s) co-terminating with a mild foot-shock (US; 0.5 mA, 2 s), repeated 6 times at 2 mins intervals.

During recall (day 2), mice were put in the same chamber with a different context (striped walls cleaned with 0.5% hydrogen peroxide). Mice were re-exposed to CS+ without foot shocks for 6 repetitions with a 2-minute inter-tone interval.

Fear learning was quantified by the percentage of time spent freezing to CS+/US during the conditioning phase, and by the percentage of time freezing to CS+ following presentation in the memory recall phase.

### Fiber photometry

A Doric system (*Doric Lenses, QC*) was used to collect all photometry data. The system consisted of a 6 port Doric Fluorescence Minicube (*IlFMC2, Doric Lenses*) equipped with 3 LEDs (405nm, 470nm, and 560nm for excitation of isosbestic-point GCaMP8f, GCaMP8f, and jRGECO1a, respectively) with integrated filters and photodetector, LED drivers (*Doric Lenses*) and a fiber photometry console (*Doric Lenses*) controlled by a computer using the Doric Neuroscience Studio software. To separate channels acquisitions, light intensity was modulated sinusoidally at non-overlapping frequencies (208.616Hz, 572.205Hz, and 333.786Hz for 405nm, 470nm, and 560nm LEDs respectively). To minimize photobleaching, light power was set to 30μW at the tip of the fiber for all LED channels. Optic fiber cannulas were connected to a low-autofluorescence patch cord (Doric Lenses, 400μm Ø) through a 2.5mm zirconia mating sleeve (Doric Lenses). Emitted light was filtered (460-490nm, 500-540nm, and 580-680nm for isosbestic-point GCaMP8f, GCaMP8f, and jRGECO1a respectively) and then collected by the integrated femtowatt photodetector and amplified 10 times (DC amplification) when necessary.

To accurately synchronize behavioral and fiber photometry data, recordings were started with anymaze via TTL pulses using anymaze synchronization mode with a 6-port anymaze synchronization interface (#60069).

### Analysis

#### Behavior analysis

All video tracking, fiber photometry synchronization, and behavior analyses were performed using AnyMaze (v7.49, Stoelting co.). Videos were recorded at 30 fps.

#### Fiber photometry analysis

All raw traces from each channel were first resampled at 60Hz and removed from recording artifacts. Photobleaching was corrected by fitting a double exponential to each trace using the least square methods, and then subtracting it from the traces. Motion correction was performed by fitting the 470nm and 560nm channels to the 405nm isosbestic channel using linear regression, then subtracting it from the 470 and 560 channels. Traces were then z-scored over the entire recording by subtracting signal at each time point from the mean signal and then dividing it by the standard deviation of the recording session.

For signal heatmap over mice position in the maze, z-scored signals were averaged over 0.5×0.5cm spatial bins and normalized. Heat maps were then smoothed by convolution with a 2D Gaussian kernel.

As all recordings were sampled at the same frequency, area under the curve per second was computed as the cumulative sum of z-scored signal during specified maze compartment, divided by time spent in specified compartment. To be included in the analysis, a compartment had to be explored by the mouse for a minimum of 5s.

For transitions between maze compartments perievent time histograms, transitions were defined as epochs in which mice spent 80% of time of the pre-defined pre-transition window in the pre-compartment before crossing and 80% of time in the pre-defined post-transition window in the post-transition compartment. PETH were then averaged for each mouse and normalized by subtracting mean of the pre-transition window.

#### Logistic Regression Classification

To assess robustness and specificity of astrocytic calcium signals during anxiogenic exploration, a logistic regression classifier was trained to predict behavior from BLA astrocytes Ca^2+^.

Data were obtained from individual mice and preprocessed to label exploration states: time spent in the closed arms of the EPM or the black alley of the SAM was coded as non-exploration (negative class, 0), while time spent outside these regions was coded as exploration (positive class, 1). Each dataset was then segmented in 5s epochs (0.5s sliding window) and only the windows that exclusively contained either exploration or non-exploration were kept for classification.

As classes were imbalanced, training datasets were balanced by randomly under-sampling the majority class. A logistic regression model with *L*_*2*_ (ridge) regularization and coordinate descent solver was trained on either EPM or SAM data, with C hyperparameter optimized via grid search using 3-fold cross-validation and area under the receiver operating characteristic curve area under the curve (ROC-AUC) as scoring metric. The best-performing model was selected and evaluated on either SAM or EPM data (for training on EPM and SAM respectively). The procedure was repeated with 10 distinct under-sampling randomizations and performance metrics of the models from all repetitions were averaged to obtain model performance for each mouse.

To compare performance against chance, each model selected after hyperparameter optimization was then trained on a shuffled dataset (following random permutation of the labels). The procedure was repeated 50 times and results of these models were averaged to generate a null distribution of classification performance for each mouse.

Accuracy was computed as:

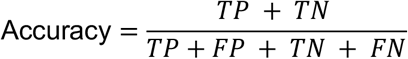

Recall was computed as:

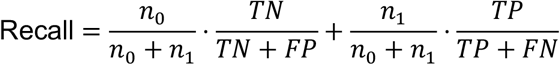

Precision was computed:

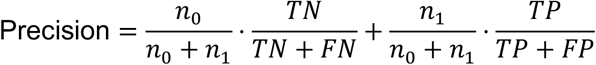

F1-score was computed as:

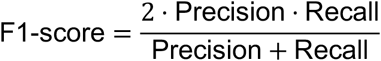

Where n_0_ is the number of negative labels, n_1_ the number of positive labels, *TN* the number of true negatives, *TP* the number of true positives, *FN* the number of false negatives, and *FP* the number of false positives.

Averaged confusion matrices were obtained by normalizing confusion matrices from each individual model before averaging.

### Immunohistochemistry

Mice were transcardially perfused with PBS, then 4% PFA. Brains were then extracted and post-fixed in 4% PFA for 24 hours at 4°C. Fixed brains were placed in a 30% sucrose solution until sank and were then flash-frozen in 2-methylbutane between –40 and –50°C and then stored at –80°C until cryosectioning in 30µm thick slices with a Leica SM 2000R microtome.

Free-floating brain slices were first washed 3 times in PBS for 15 mins. Slices were permeabilized using a block-perm solution (3% bovine serum albumin, 0.5% Triton10% in PBS) for one hour. Slices were incubated with primary antibodies in block-perm for 24 hours ([rabbit] anti-S100β 1:1000, Abcam, ab52642; [chicken] anti-eGFP 1:1000, Thermofisher, TA100022; [mouse] anti-NeuN, 1:500, Millipore Sigma, MAB377). Slices were then washed 3 times, before incubation with secondary antibodies in DAPI (1:10000 in PBS, secondary antibodies: [goat] anti-mouse Alexa 647, 1:1000, Thermofisher, A21449; [donkey] anti-chicken Alexa 488, Jackson Immuno Research, 711-545-152, 1:1000; [goat] anti-rabbit Alexa 568, 1:1000, Thermofisher, A11011). After secondary antibody incubation, slices were washed again three times in 1× PBS for 15 min before mounting onto microscope slides (Fisherbrand) using Antifade mounting medium (ProLong Glass, P36982).

### Microscopy

Images were acquired with either a Zeiss Observer Z1 spinning disk confocal microscope/TIRF with a ×20 objective 1.8 NA (viral validation) or an Olympus Optical microscope BX61VSF with a 20X 0.75NA objective and a resolution of 0.325µm (fiber implant validation). For viral validation, 20μm z-stack images (1μm step) were acquired, and 10μm maximum intensity z-stack projections were analyzed using the cell-count plugin in FIJI.

### Statistical analysis

All results are represented as mean ± SEM. Graphical significance levels were *p<0.05; **p<0.01, and ***p<0.001.

We used paired or independent Student’s t-tests for comparisons between two groups, for paired and independent groups comparison respectively. For variables including more than 2 repeated observations, we used repeated measure ANOVA with greenhouse-geisser correction, followed by Tukey’s HSD test for multiple comparisons. For analysis including 2 variables, we used linear mixed-effect models followed by Tukey’s HSD test for multiple comparisons

Normality was assessed via Shapiro tests and appropriate non-parametric alternatives to parametric tests were used when necessary.

All statistical analyses were conducted using custom Python scripts.

## Data and Code Availability

All data associated with this manuscript will be made freely accessible upon publication. Custom code for decoding analysis is available on GitHub at: https://github.com/Ossamaghenissa/anxiety-astroBLA-paper

---

**Figure S1.**
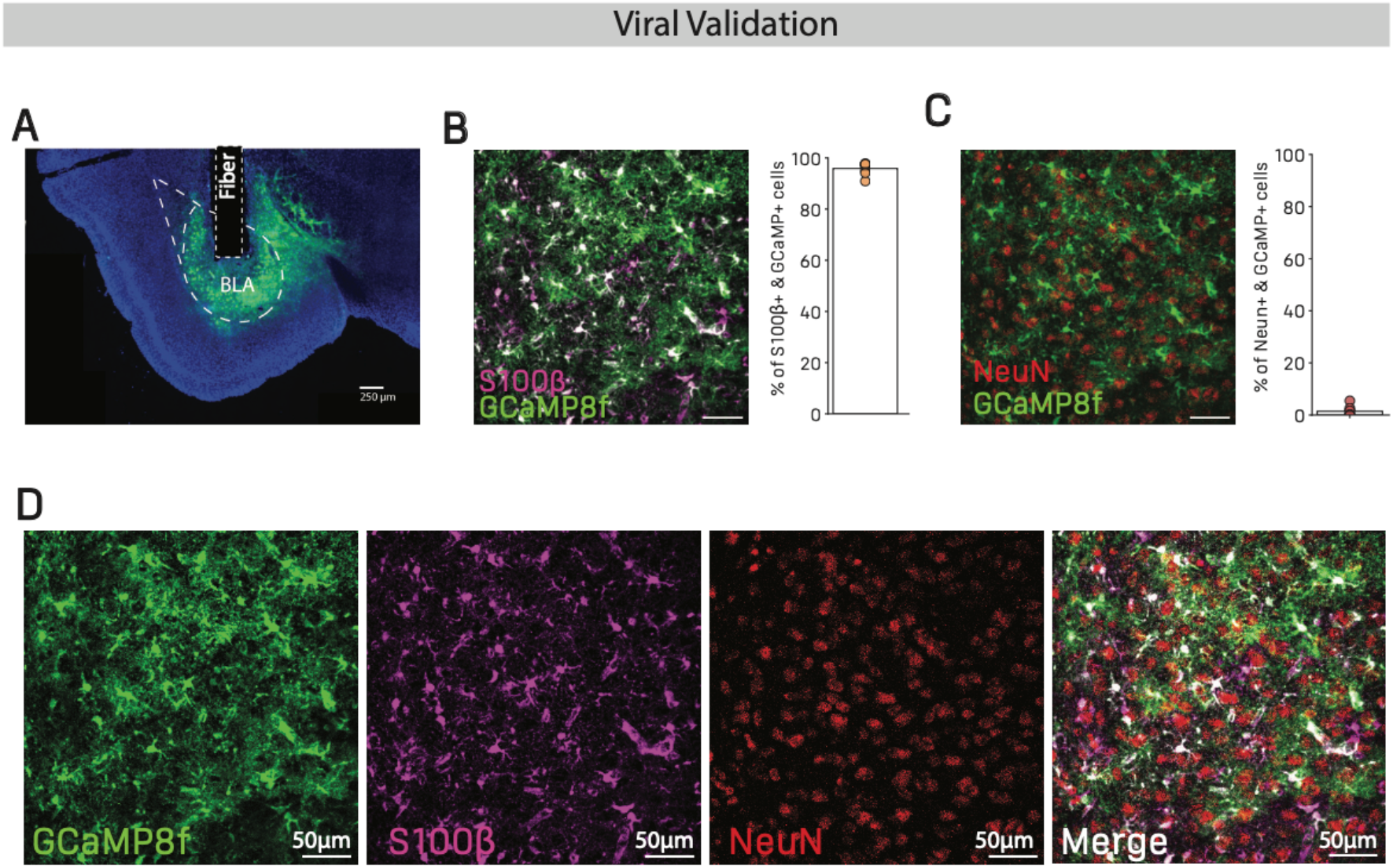
A) Representative image of injection site and optic fiber implantation, scalebar = 250µm B) Representative image for GCaMP8f-S100β colocalization, 95.89±0.85% S100β positive of GCaMP8f positive cells, n=8 slices from 6 mice. C) Representative image for GCaMP8f-NeuN colocalization, 1.42±0.68% NeuN positive of GCaMP8f positive cells, n=8 slices from 6 mice. D) Individual channels from representative image of immunostainings, from left to right (1) astrocytic GCaMP8f expression, (2) astrocyte marker S100B, (3) neuronal nuclei marker NeuN, (4) Merge.

**Figure S2.**
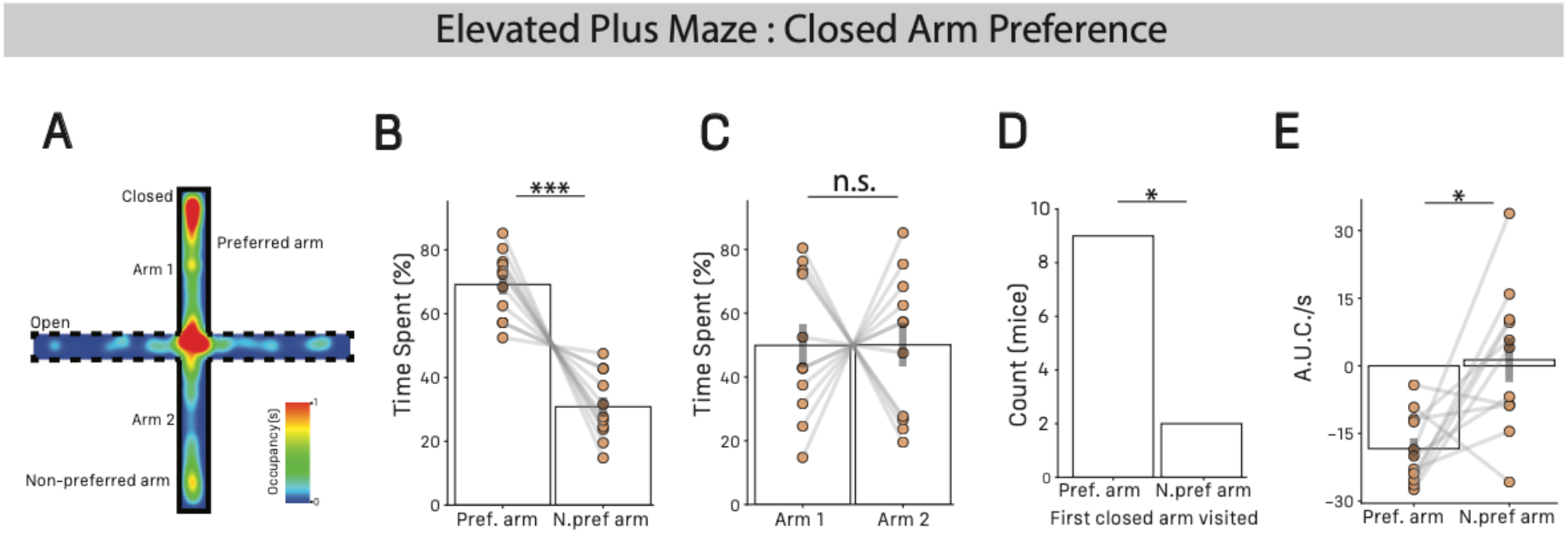
A) Representative occupancy map in the EPM. B) Time spent in preferred and non-preferred arm, preferred = 69.17±3.17, non-preferred = 30.82±3.17, paired T-test T=6.03, p<0.001, n=11 mice. C) Time spent in closed arm 1 and 2, arm 1 = 49.91±6.84, arm 2 = 50.08±6.84, paired T-test, T=0.013, p=0.98, n=11 mice. D) Count of mice that first visited the preferred vs. non-preferred arm, binomial test, p=0.032, n=11 mice. E) Astrocyte area under the curve per second for preferred and non-preferred arm, preferred = −18.37±1.33, non-preferred = 1.33±4.94, paired T-test T=-3.08, p=0.011), n=11 mice. All results are represented as mean ± SEM. Graphical significance levels are *p<0.05; **p<0.01, and ***p<0.001.

**Figure S3.**
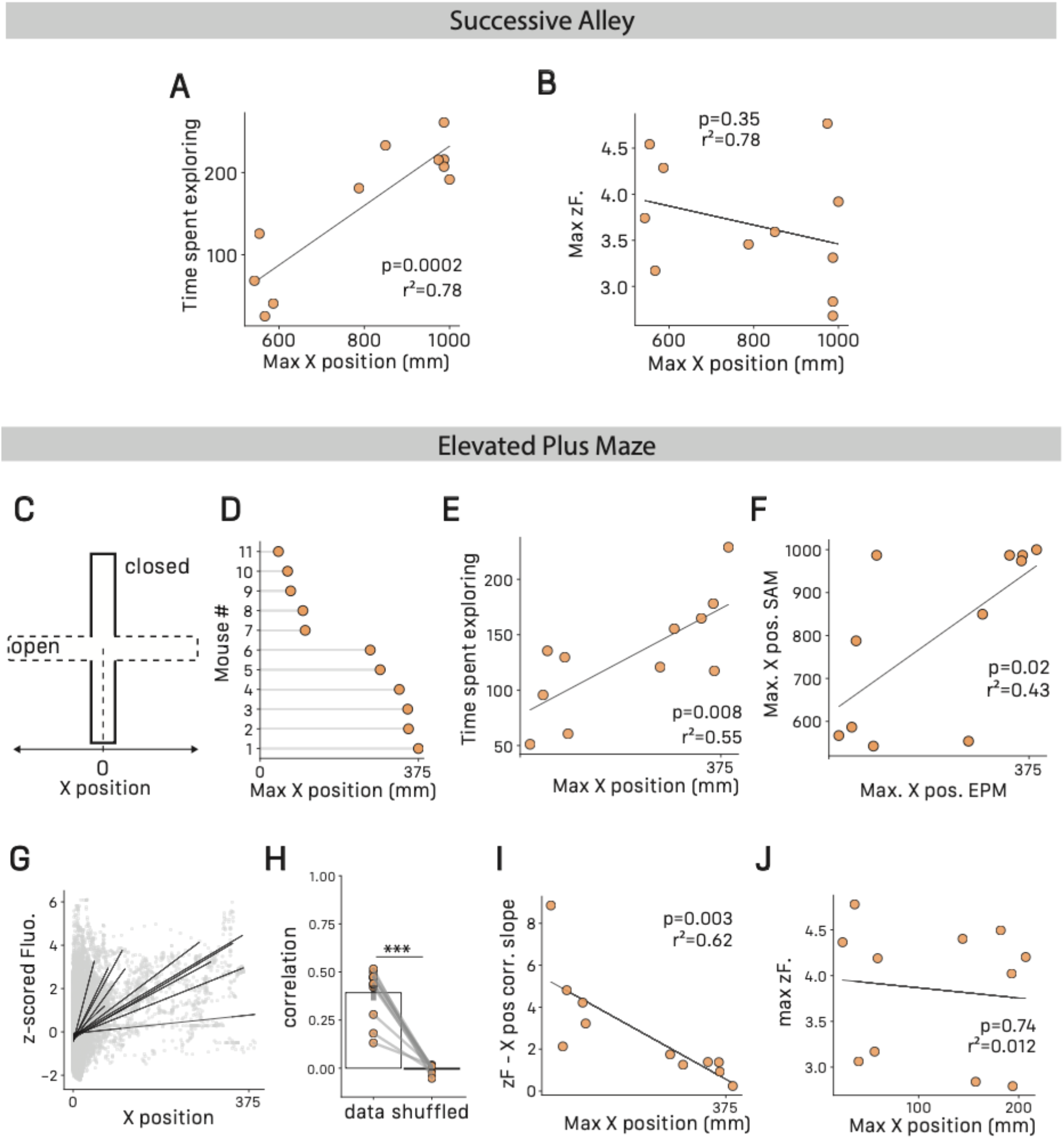
A) Maximum X position reached in the SAM through time spent exploring anxiogenic compartments of the maze (alleys 2, 3, or 4). Linear regression, r^2^=0.78, n=11 mice. B) Maximum X position reached in the SAM through maximum astrocyte z-scored fluorescence (average of 99^th^ quantile), linear regression, r^2^=0.095, p=0.35, n=11 mice. C) Schematic of X position along the open arm axis of the EPM. D) Maximum X position in the EPM for individual mice, n=11 mice. E) Maximum X position reached in the EPM through time spent exploring anxiogenic compartments in the EPM (center or open arms), linear regression r^2^=0.55, p<0.001, n=11 mice. F) Maximum X position reached in the EPM through maximum X position reached in the SAM, linear regression, r^2^=0.43, p=0.02, n=11 mice. G) Correlation between X position and astrocytes z-scored Fluorescence, black lines indicate individual regression lines, n=11 mice. H) Coefficients from X position-astrocyte z-score correlation for true data and shuffled dataset, data=0.39±0.04, shuffled=-0.005±0.005, n=11 mice. I) Maximum X position reached in the EPM through astrocyte z-scored fluorescence, linear regression, r^2^=0.62, p=0.003, n=11 mice. J) Maximum X position reached in the EPM through maximum astrocyte z-scored fluorescence (average of values above 99^th^ quantile), linear regression, r^2^=0.012, p=0.74, n=11 mice. All results are represented as mean ± SEM. Graphical significance levels are *p<0.05; **p<0.01, and ***p<0.001.

**Figure S4.**
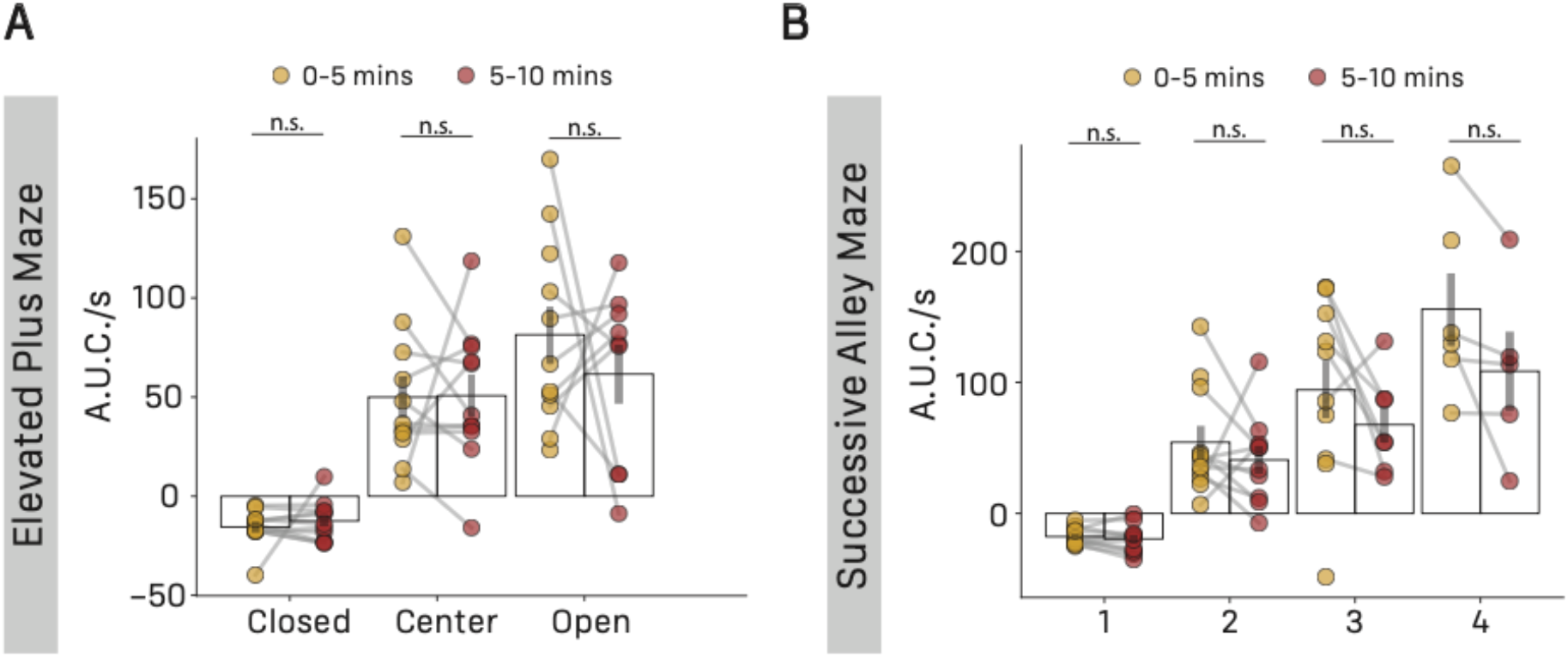
A) Astrocytes area under the curve per second through SAM compartments for first and second half of recording, 2-way ANOVA, main effect of recording period F_1,4_=2.93, p=0.16, n=11 mice. B) Astrocytes area under the curve per second through SAM compartments for first and second half of recording, 2-way ANOVA, main effect of recording period F_1,8_=0.43, p=0.52, n=11 mice. All results are represented as mean ± SEM. Graphical significance levels are *p<0.05; **p<0.01, and ***p<0.001.

**Figure S5.**
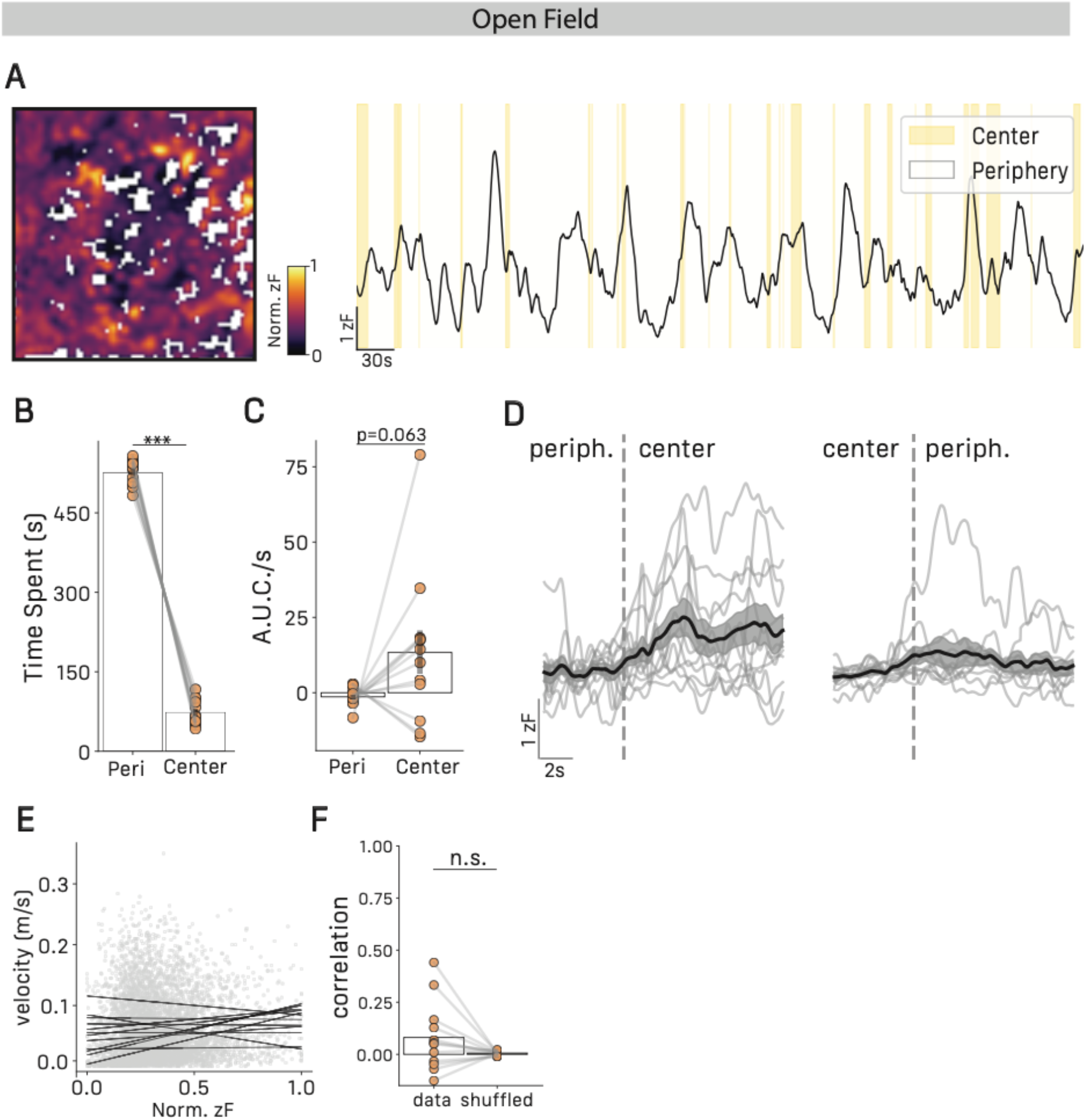
A) Representative heat map (left) and trace (right) of astrocyte Ca2+ activity in the open field. B) Proportion of time spent in open field compartments, periphery=87.44±1.25, center=12.16±1.25, paired T-test T=29.98, p<0.001, n=11 mice. C) Astrocytes area under the curve per second in open field compartments, periphery=- 1.35±0.85, center=13.37±7.31, Wilcoxon signed-ranks test W=15, p=0.063, n=11 mice. D) Average z-scored fluorescence traces of astrocyte activity during periphery exits (left) and entries (right), n=11, gray traces represent individual averages, n=11 mice. E) Normalized astrocytes z-scored fluorescence through velocity in the open field, black lines indicate regression lines for individuals, n=11 mice. F) Coefficients from astrocytes z-score-velocity correlation for true data and shuffled data, data=0.08±0.04, shuffled=0.006±0.002, paired T-test, T=23, p<0.23, n=11 mice. All results are represented as mean ± SEM. Graphical significance levels are *p<0.05; **p<0.01, and ***p<0.001.

**Figure S6.**
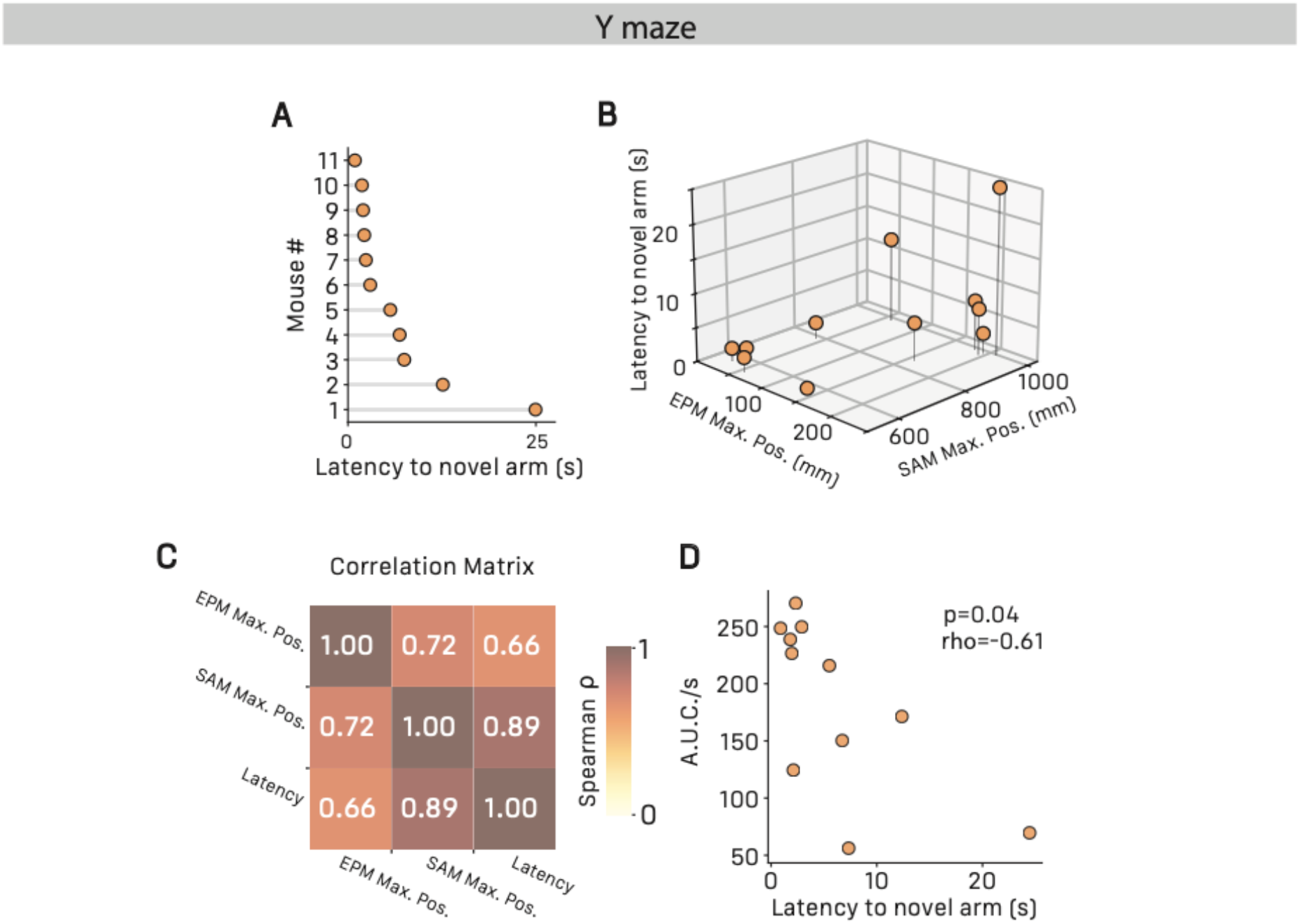
A) Latency to enter novel arm in the Y-maze novel arm task, n=11 mice. B) Latency to enter novel arm in the Y-maze task through maximum X position reached in the EPM and maximum X position reached in the SAM, n=11 mice. C) Correlation matrix for latency to enter novel arm in the Y-maze task, maximum X position reached in the EPM and maximum X position reached in the SAM, Spearman correlations, Y-maze-EPM, ρ=0.66, p=0.02; Y-maze-SAM, ρ=0.88, p<0.001; EPM-SAM, ρ=0.72, p=0.01, n=11 mice. D) Correlation between latency to enter novel arm and astrocyte area under the curve per second during first entry in the novel arm in the Y-maze novel arm task, spearman correlation, ρ=-0.61, p=0.04, n=11 mice. All results are represented as mean ± SEM. Graphical significance levels are *p<0.05; **p<0.01, and ***p<0.001.

**Figure S7.**
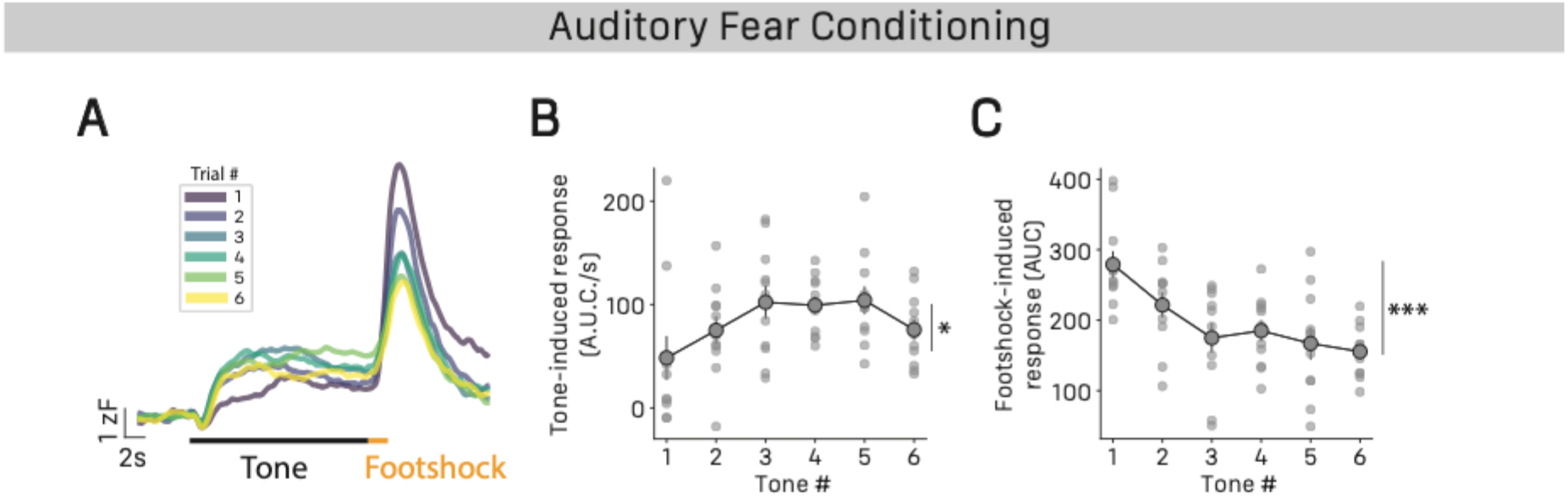
A) Average astrocyte z-scored fluorescence traces through trials during acquisition phase of auditory fear conditioning, n=11 mice. B) Astrocytes area under the curve per second through tone number during acquisition phase, repeated measure ANOVA, effect of tone number F5_,50_=2.94, p=0.02, n=11mice. C) Astrocytes area under the curve per second through foot-shock number presentation during acquisition phase, repeated measure ANOVA, effect of foot-shock number F5_,50_=2.94, p<0.001, n=11mice. All results are represented as mean ± SEM. Graphical significance levels are *p<0.05; **p<0.01, and ***p<0.001.

**Figure S8.**
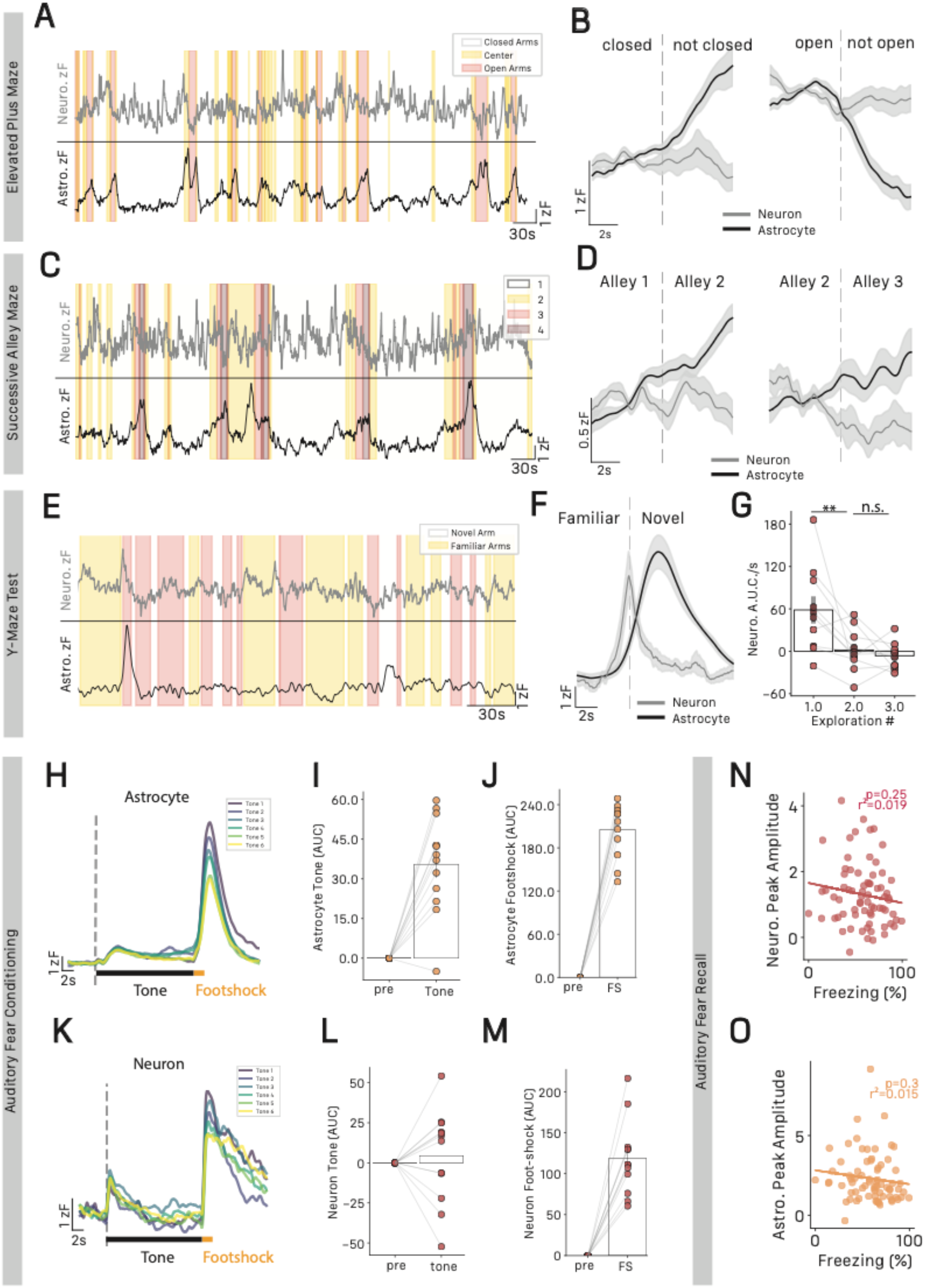
A. Representative trace of dual recordings of neurons (top) and astrocytes (bottom) during EPM. B. Average z-scored fluorescence traces of astrocytes and neuronal activity during closed arms exits (left) and entries (right), n=12 mice. C. Representative trace of dual recordings of neurons (top) and astrocytes (bottom) during SAM. D. Average z-scored fluorescence traces of astrocytes and neuronal activity during transition from alley 1 to 2 (left) and 2 to 3 (right), n=12 mice. E. Representative trace of dual recordings of neurons (top) and astrocytes (bottom) during the Y-maze novel arm task. F. Average z-scored fluorescence traces of astrocytes and neuronal activity during first exploration of the novel arm, n=12 mice. G. Neurons area under the curve per second for successive bouts of exploration of the novel arm. 1^st^ bout = 68.5±19.29, 2^nd^ bout = 1.171±9.59, 3^rd^ bout = −6.37±5.93, repeated measure ANOVA F_2,18_=8.15, p=0.003, Tukey HSD 1^st^ vs. 2^nd^ p=0.017, 2^nd^ vs. 3^rd^ p=0.89, n=12 mice. H. Average astrocyte z-scored fluorescence traces through trials during acquisition phase of auditory fear conditioning, n=12 mice. I. Astrocytes area under the curve per second for pre-tone vs. tone presentation, pre-tone=0±0, tone=35.47±5.32, paired T-test, T=6.66, p<0.001, n=12 mice. J. Astrocytes area under the curve per second for pre-shock vs. foot-shock presentation, pre-tone=0±0, tone=205.33±10.9, paired T-test, T=18.83, p<0.001, n=12 mice. K. Average neuron z-scored fluorescence traces through trials during acquisition phase of auditory fear conditioning, n=12 mice. L. Neuron area under the curve per second for pre-tone vs. tone presentation, pre-tone=0±0, tone=4.44±8.48, paired T-test, T=0.52, p=0.6, n=12 mice. M. Neuron area under the curve per second for pre-shock vs. foot-shock presentation, pre-tone=0±0, tone=205.33±10.9, paired T-test, T=18.83, p<0.001, n=12 mice. N. Proportion of time spent freezing and neuronal peak amplitude during recall phase, linear regression, r^2^=0.019, p=0.25, n=12 mice. O. Proportion of time spent freezing and astrocytic peak amplitude during recall phase, linear regression, r^2^=0.019, p=0.25, n=12 mice. All results are represented as mean ± SEM. Graphical significance levels are *p<0.05; **p<0.01, and ***p<0.001.

**Figure S9.**
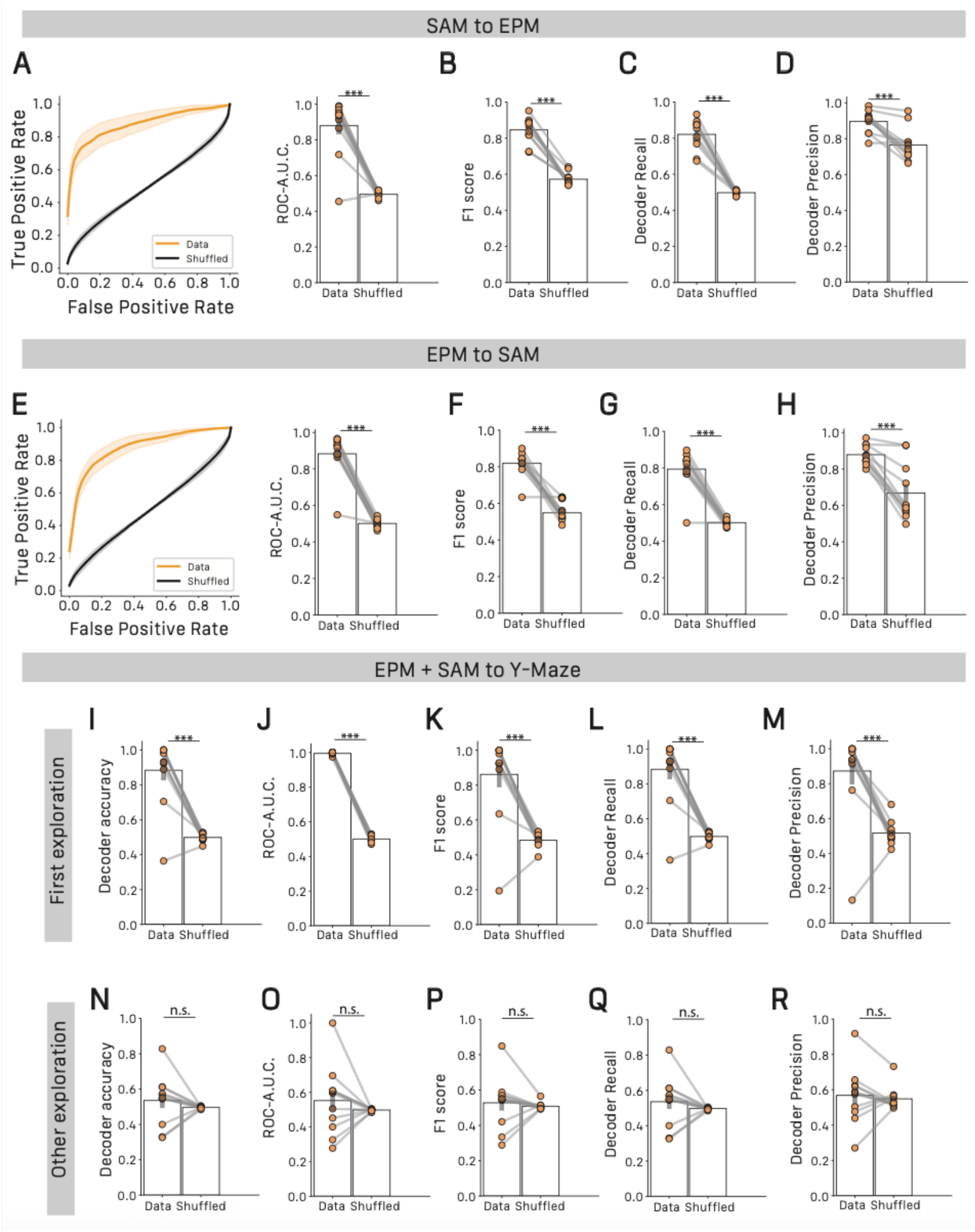
A. Receiver operator characteristic (ROC) curves (left) and ROC-area under the curve for SAM-to-EPM true data and shuffled labels decoder, true data=0.88±0.035, shuffled=0.5±0.007, paired T-test, T=9.912, p<0.001, n=11 mice. B. F1 score for SAM-to-EPM true data and shuffled labels decoder, true data=0.81±0.02, shuffled=0.548±0.014, Paired T-test T=10.7, p<0.001, n=11 mice. C. Recall score for SAM-to-EPM true data and shuffled labels decoder, true data=0.82±0.02, shuffled=0.497±0.004, Paired T-test T=12.63, p<0.001, n=11 mice. D. Precision score for SAM-to-EPM true data and shuffled labels decoder, true data=0.896±0.018, shuffled=0.765±0.028, Paired T-test T=3.89, p<0.001, n=11 mice. E. ROC curves (left) and ROC-area under the curve for EPM-to-SAM true data and shuffled labels decoder, true data=0.87±0.04, shuffled=0.495±0.005, paired T-test, T=10.65, p<0.001, n=11 mice. F. F1 score for EPM-to-SAM true data and shuffled labels decoder, true data=0.81±0.02, shuffled=0.54±0.014, Paired T-test T=10.7, p<0.001, n=11 mice. G. Recall score for EPM-to-SAM true data and shuffled labels decoder, true data=0.82±0.02, shuffled=0.497±0.004, Paired T-test T=12.63, p<0.001, n=11 mice. H. Precision score for EPM-to-SAM true data and shuffled labels decoder, true data=0.878±0.016, shuffled=0.667±0.046, Paired T-test T=4.27, p<0.001, n=11 mice. I. Accuracy score for EPM+SAM-to-Y maze true data and shuffled labels decoder on first exploration, true data=0.88±0.058, shuffled=0.49±0.006, paired T-test, T=6.58, p<0.001, n=11 mice. J. ROC-area under the curve for SAM-to-EPM true data and shuffled labels decoder on first exploration, true data=0.994±0.002, shuffled=0.5±0.006, paired T-test, T=74.58, p<0.001, n=11 mice. K. F1 score for EPM-to-SAM true data and shuffled labels decoder on first exploration, true data=0.861±0.07, shuffled=0.483±0.014, Paired T-test T=6.23, p<0.001, n=11 mice. L. Recall score for EPM-to-SAM true data and shuffled labels decoder on first exploration, true data=0.883±0.058, shuffled=0.498±0.006, Paired T-test T=6.58, p<0.001, n=11 mice. M. Precision score for EPM-to-SAM true data and shuffled labels decoder on first exploration, true data=0.872±0.076, shuffled=0.516±0.02, Paired T-test T=4.47, p<0.001, n=11 mice. N. Accuracy score for EPM+SAM-to-Y maze true data and shuffled labels decoder on subsequent exploration, true data=0.53±0.043, shuffled=0.497±0.001, paired T-test, T=0.89, p=0.38, n=11 mice. O. ROC-area under the curve for SAM-to-EPM true data and shuffled labels decoder on subsequent exploration, true data=0.552±0.059 shuffled=0.497±0.001, paired T-test, T=0.912, p<0.001, n=11 mice. P. F1 score for EPM-to-SAM true data and shuffled labels decoder on subsequent exploration, true data=0.527±0.044, shuffled=0.508±0.006, Paired T-test T=0.43, p=0.671, n=11 mice. Q. Recall score for EPM-to-SAM true data and shuffled labels decoder on subsequent exploration, true data=0.535±0.043, shuffled=0.497±0.001, Paired T-test T=0.89, p=0.38, n=11 mice. R. Precision score for EPM-to-SAM true data and shuffled labels decoder on subsequent, true data=0.568±0.048, shuffled=0.548±0.019, Paired T-test T=0.383, p=0.7, n=11 mice. All results are represented as mean ± SEM. Graphical significance levels are *p<0.05; **p<0.01, and ***p<0.001. # indicates significance against shuffled labels decoder equivalent.

**Figure S10.**
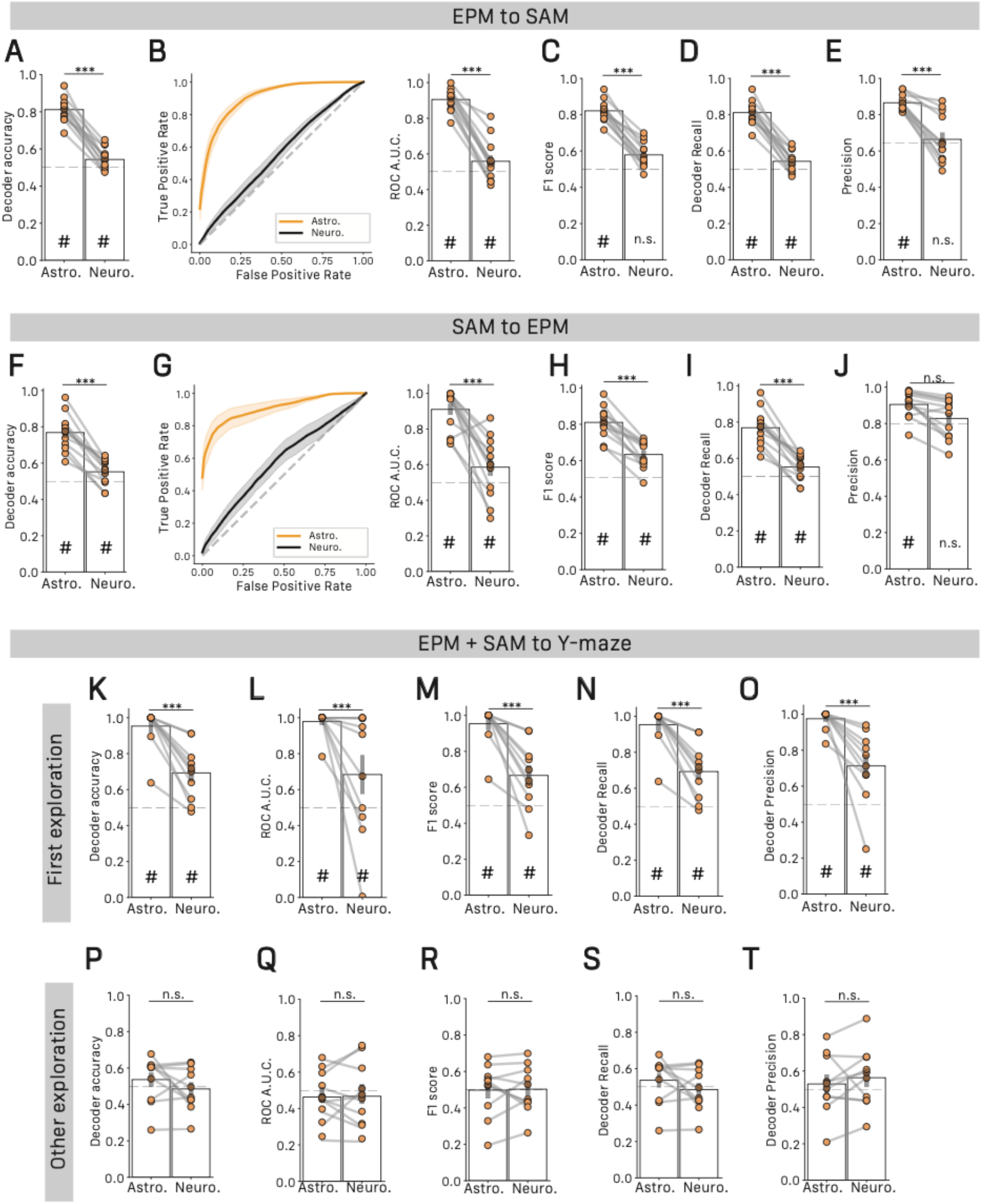
A) Accuracy score for EPM-to-SAM astrocytes and neuronal decoder, astrocytes=0.811±0.018, neurons=0.543±0.016, paired T-test, T=10.65, p<0.001, n=12 mice. B) Receiver operator characteristic (ROC) curves (left) and ROC-area under the curve for EPM-to-SAM astrocytes and neuronal decoder, astrocytes=0.90±0.017, neuronal=0.558±0.033, paired T-test, T=9.04, p<0.001, n=12 mice. C) F1 score for EPM-to-SAM astrocytes and neuronal decoder, astrocytes=0.821±0.016, neurons=0.578±0.02, paired T-test, T=9.15, p<0.001, n=12 mice. D) Recall score for EPM-to-SAM astrocytes and neuronal decoder, astrocytes=0.811±0.018, neurons=0.543±0.016, paired T-test, T=10.65, p<0.001, n=12 mice. E) Precision score for EPM-to-SAM astrocytes and neuronal decoder, astrocytes=0.864±0.012, neurons=0.664±0.038, paired T-test, T=4.95, p<0.001, n=12 mice. F) Accuracy score for SAM-to-EPM astrocytes and neuronal decoder, astrocytes=0.768±0.029, neurons=0.552±0.02, paired T-test, T=6.07, p<0.001, n=12 mice. G) Receiver operator characteristic (ROC) curves (left) and ROC-area under the curve for SAM-to-EPM astrocytes and neuronal decoder, astrocytes=0.91±0.033, neuronal=0.558±0.049, paired T-test, T=5.44, p<0.001, n=12 mice. H) F1 score for SAM-to-EPM astrocytes and neuronal decoder, astrocytes=0.81±0.024, neurons=0.633±0.021, paired T-test, T=5.35, p<0.001, n=12 mice. I) Recall score for SAM-to-EPM astrocytes and neuronal decoder, astrocytes=0.768±0.029, neurons=0.55±0.02, paired T-test, T=6.07, p<0.001, n=12 mice. J) Precision score for SAM-to-EPM astrocytes and neuronal decoder, astrocytes=0.904±0.02, neurons=0.822±0.031, paired T-test, T=2.05, p=0.051, n=12 mice. K) Accuracy score for EPM+SAM-to-Y maze astrocytes and neuronal decoder on first exploration, astrocytes=0.953±0.046, neurons=0.69±0.048, paired T-test, T=4.27, p<0.001, n=12 mice. L) ROC-area under the curve for EPM+SAM-to-Y maze astrocytes and neuronal decoder on first exploration, astrocytes=0.978±0.109, neurons=0.68±0.109, paired T-test, T=2.64, p<0.001, n=12 mice. M) F1 score for EPM+SAM-to-Y maze astrocytes and neuronal decoder on first exploration, astrocytes=0.953±0.036, neurons=0.665±0.058, paired T-test, T=4.18, p<0.001, n=12 mice. N) Recall score for EPM+SAM-to-Y maze astrocytes and neuronal decoder on first exploration, astrocytes=0.953±0.046, neurons=0.69±0.048, paired T-test, T=4.27, p<0.001, n=12 mice. O) Precision score for EPM+SAM-to-Y maze astrocytes and neuronal decoder on first exploration, astrocytes=0.975±0.017, neurons=0.712±0.064, paired T-test, T=3.94, p<0.001, n=12 mice. P) Accuracy score for EPM+SAM-to-Y maze astrocytes and neuronal decoder on subsequent exploration, astrocytes=0.536 ±0.04, neurons=0.485±0.037, paired T-test, T=0.92, p=0.36, n=12 mice. Q) ROC-area under the curve for EPM+SAM-to-Y maze astrocytes and neuronal decoder on subsequent explorations, astrocytes=0.462±0.044, neurons=0.467±0.057, paired T-test, T=0.06, p=94, n=12 mice. R) F1 score for EPM+SAM-to-Y maze astrocytes and neuronal decoder on subsequent exploration, astrocytes=0.497 ±0.05, neurons=0.502±0.041, paired T-test, T=0.08, p=0.93, n=12 mice. S) Recall score for EPM+SAM-to-Y maze astrocytes and neuronal decoder on subsequent exploration, astrocytes=0.536 ±0.04, neurons=0.485±0.037, paired T-test, T=0.92, p=0.36, n=12 mice. T) Precision score for EPM+SAM-to-Y maze astrocytes and neuronal decoder on subsequent exploration, astrocytes=0.528 ±0.052, neurons=0.563±0.052, paired T-test, T=0.46, p=0.65, n=12 mice. All results are represented as mean ± SEM. Graphical significance levels are *p<0.05; **p<0.01, and ***p<0.001. # indicates significance against shuffled labels decoder equivalent.

